# Genomic footprints of domestication in almond (*Prunus dulcis*)

**DOI:** 10.1101/2025.08.06.668913

**Authors:** C. Lougmani, A. Mesnil, R. Fahmy, V. Decroocq, Q. T. Bui, S. Liu, X. Chen, A. Chague, A. Venon, D. Chagné, I. Eduardo, K. Alix, A. Cornille

**Author notes:** Contributed equally (C. Lougmani, A. Mesnil).

## Abstract

The domestication of perennial crops in the Mediterranean Basin remains unclear, particularly regarding the genomic consequences of human-mediated demographic shifts and selection. We analyzed 8.1 million single nucleotide polymorphisms from 96 cultivated almond (*Prunus dulcis*) accessions from Europe, North America, Central Asia, and New Zealand, alongside four wild relatives. Population structure analyses revealed four geographically differentiated cultivated groups (Central Asian, North American, and two European) and three wild populations (*P. spinosissima*, *P. orientalis*, and *P. fenzliana*). Cultivated almonds retained high genetic diversity, consistent with weak domestication bottlenecks typical of outcrossing perennials. Elevated diversity and private allele counts in Central Asian cultivars, with limited gene flow, support this region as an independent cradle of domestication. In contrast, extensive wild-to-crop gene flow—especially involving *P. orientalis*—has shaped the genomes of European and North American almonds. Genome-wide scans for selective sweeps showed most candidate genes under selection were population-specific, though often associated with similar biological functions, including stress responses and agronomic traits. This suggests repeated targeting of comparable pathways during domestication, despite distinct selection histories. Several selected genes in cultivated populations overlapped with those in wild relatives, particularly *P. orientalis*. Combined with demographic inferences indicating wild populations persistence through past climate fluctuations, these findings suggest wild gene pools retain adaptive alleles—either ancestrally shared or introgressed—that contributed to cultivated diversity. Altogether, our results reveal a complex, multi-regional domestication history for almonds, shaped by gene flow and recurrent selection. This study emphasizes wild relatives as adaptive diversity sources and reveals genomic bases of perennial crop evolution.

## Introduction

Plant domestication began approximately 10,000 years ago, at the onset of the Holocene epoch, when some human societies transitioned from a hunter-gatherer lifestyle to a sedentary one fueled by the emergence of agriculture (Brown et al., 2009; Meyer et al., 2012). Humans began to selectively cultivate plants with desirable traits, such as larger fruits with lower toxicity, non-shattering seed pods, pest resistance, and limited branching (Dong et al., 2019; Glémin & Bataillon, 2009; Sánchez-Pérez et al., 2019; Tsiantis, 2011; Wang et al., 2023), leading to the emergence of domesticated crop taxa that are genetically and morphologically distinct from their wild counterparts (Gaut et al., 2015; Glémin & Bataillon, 2009; Meyer et al., 2012; Purugganan, 2019; Zohary et al., 2012). As it drives the emergence of new varieties, populations, and species, thereby contributing to biodiversity (Darwin, 1859; Gaut et al., 2015; Glémin & Bataillon, 2009; Purugganan, 2019), how domestication has shaped the genomes of modern crops has been a key topic in ecology and evolution. Research focus in this field addresses the origins and progenitors of domesticated species, the diffusion of crops following their domestication, population structure, and genetic diversity among wild and domesticated taxa, demographic variation within crops, the extent of gene flow between wild and cultivated populations, and the identification of genes under selection during domestication. Beyond its academic significance, understanding domestication is also crucial for advancing crop breeding programs, as it provides insights into the genomic bases of important agronomic traits (Ross-Ibarra et al., 2007). In this context, the wild relatives of crops represent a largely untapped reservoir of genetic diversity, offering immense potential for crop improvement (Dempewolf et al., 2017). Despite widespread interest in domestication from a historical perspective, many aspects of the evolutionary trajectory and genomic basis that have led to the differentiation of crops from their wild relatives remain under-investigated, particularly in perennial crops.

Domestication of perennial crops has received less research attention than that of annual crops (Gaut et al., 2015). Differences in life-history traits, such as longer generation times, extended juvenile phases, and higher rates of outcrossing due to self-incompatibility systems, make perennials fundamentally different from annuals in their domestication trajectories (Gaut et al., 2015). Studies on perennial crop domestication have largely focused on fruit trees, which are keystone species in forests and agroecosystems. Among perennial fruit trees, the domestication of Rosaceae family members began approximately 4,000 years ago (Zohary & Spiegel-Roy, 1975), about 6,000 years later than with cereals and coinciding with the emergence of vegetative propagation techniques such as cuttings and grafting, which facilitated the cultivation of many fruit tree species (Meyer et al., 2012; Zohary & Spiegel-Roy, 1975). As a result, fewer generations have elapsed since the domestication of perennial species compared to annual species, leading to a lower degree of genetic differentiation from their wild progenitors and making domestication studies more challenging for fruit trees (Gaut et al., 2015). Genomic studies have confirmed the protracted nature of crop domestication (Allaby et al., 2008; Brown et al., 2009; Gaut et al., 2015; Purugganan, 2019). For species such as sweet cherries (*Prunus avium*), pears (*Pyrus* spp.), and lychees (*Litchi chinensis*), domestication occurred multiple times across different geographical regions and over extended periods (Hu et al., 2022; Mariette et al., 2010). A recent discovery in apricots (*Prunus armeniaca*) revealed two independent domestication events from distinct Central Asian wild populations, leading to the formation of two differentiated gene pools—the European and Chinese apricots—connected by subsequent recent hybridizations in the 20^th^ century (Groppi et al., 2021; Liu et al., 2019). The cultivated apple (*M. domestica*) originated from a domestication event in Central Asia, followed by introgressions from several wild species along the Silk Road (Bina et al., 2022; Chen et al., 2023; Cornille et al., 2012, 2014), and a second center of domestication has recently been identified in Iran (Bina et al., 2022). These studies highlight the role of gene flow in perennial species domestication, as many fruit trees exhibit allogamy—with a few exceptions, such as peach (*Prunus persica*) (Velasco et al., 2016)—leading to higher gene flow and weaker domestication bottlenecks compared to annual species (Gaut et al., 2015). For instance, gene flow from wild relatives to cultivated varieties has been observed in sweet cherry, apple, grapevine (*Vitis vinifera*), and Chinese apricot (Bina et al., 2022; Chen et al., 2023; Cornille et al., 2012; Dong et al., 2023; Freitas et al., 2021; Groppi et al., 2021; Mariette et al., 2010). High levels of gene flow have helped maintain genetic diversity in perennial crops (Gaut et al., 2015; Halász et al., 2019), and in some cases, such as cherries, apples, and Chinese common apricots, cultivated populations exhibit levels of genetic diversity comparable to or even higher than those in their wild counterparts (Cornille et al., 2012; Groppi et al., 2021; Mariette et al., 2010).

By contrast, many annual crop species have undergone severe domestication bottlenecks, resulting in a substantial loss of genetic diversity (Glémin & Bataillon, 2009; Meyer et al., 2012). Domestication also involves the selection of alleles/genes underlying important agronomic traits, either from standing genetic variation or new mutations. Bottom-up approaches, which identify genomic signatures of selection before linking them to phenotypes, are effective in studying domestication (Ross-Ibarra et al., 2007). These methods rely on molecular markers such as single-nucleotide polymorphisms (SNPs) to detect selection signals while accounting for the domestication history of the crop in question to minimize false positives (Tiffin & Ross-Ibarra, 2014). Despite growing interest in crop domestication, our understanding of the genomic basis of domestication in long-lived perennial fruit trees remains limited. Compared to annual crops, perennial species such as apricots, apples, and peaches have received less attention, and several key model systems are still under-investigated.

The cultivated almond (*Prunus dulcis* Mill. DA Webb) is a symbol of the Mediterranean Basin and a significant nut crop worldwide. In 2021, worldwide almond production exceeded three million tons across 46 countries, with the majority concentrated in California, the Mediterranean Basin, and Asia (FAO, 2023; Mahhou & Dennis, 1992). Despite the economic and agricultural importance of almond trees and growing research efforts, significant uncertainties remain regarding the domestication history and diffusion of almonds. Most genetic studies of almonds have relied on simple sequence repeat (SSR) markers or a few SNPs (∼5000) (Decroocq et al., 2025; Delplancke et al., 2016; Fernández I Martí et al., 2009; Halász et al., 2019; Rubio-Cabetas et al., 2024; Velasco et al., 2016; Wang et al., 2021; Zeinalabedini et al., 2010), often focusing on a narrow representation of diversity in cultivated almonds (Delplancke et al., 2013; Velasco et al., 2016) and frequently excluding wild almond relatives (Velasco et al., 2016). Population genetics studies suggest that initial almond domestication may have occurred approximately 5,000 years ago (Zohary et al., 2012) in Central or Western Asia without a severe domestication bottleneck (Delplancke et al., 2013; Halász et al., 2019; Velasco et al., 2016), and was followed by a separate domestication event on the eastern side of the Mediterranean Basin (Browicz & Zohary, 1996; Delplancke et al., 2013; Rahemi et al., 2012; Wang et al., 2021). The genetic clustering of cultivated almond varieties, with each regional group (e.g., Central Asian, North American) forming distinct clusters based on highly evolving neutral genetic markers (Fernández I Martí et al., 2009; Halász et al., 2019), suggests distinct evolutionary trajectories across different regions. This observation also indicates that cultivated almonds may have originated from multiple domestication events, possibly involving different wild ancestors, with recurrent gene flow from wild to crops. However, no conclusive genetic evidence has been obtained to confirm the contributions of various wild species as progenitors or through introgression. More than 30 wild almond species exist, any of which might have contributed to the genetic makeup of cultivated almonds (Browicz & Zohary, 1996; Wang et al., 2021). Among them, *Prunus fenzliana* Fritsch, native to Northeast Iran, has been suggested as a potential ancestor of *P. dulcis* (Zeinalabedini et al., 2010). Recent SSR studies indicate that *Prunus orientalis* (Mill.) Koehne, a wild almond species from East Anatolia, is genetically closer to *P. dulcis* than *P. fenzliana* to *P. dulcis*, suggesting that *P. orientalis* may be the primary ancestor or at least a significant genomic contributor to cultivated almonds (Decroocq et al., 2025). Other wild relatives, such as *Prunus turcomanica* (Lincz.) Kitam from Central Asia and *Prunus spinosissima* (Bunge) Franch., remain understudied. The occurrence of naturalized *P. dulcis* populations outside its center of origin and main cultivation areas may obscure its evolutionary history and complicate the identification of its primary wild progenitors. In New Zealand, for example, back-to-wild almonds have become invasive after being introduced by European settlers in the 19th century and subsequently abandoned, along with gold mines; however, the ancestry of these populations remains unknown. Thus, the evolutionary history of the cultivated almond, including its domestication events and global dispersal, remains incompletely understood. Likewise, the number and nature of genes selected during almond domestication remain largely unexplored. Almonds are cultivated primarily for their edible sweet kernel, for oil production, and their ornamental value (Covert, 2011). Major domestication traits include a lower toxicity and bitterness due to cyanogenic compounds (Sánchez-Pérez et al., 2019), a thinner endocarp (Goonetilleke et al., 2023; Pavan et al., 2021; Pérez De Los Cobos et al., 2021; Sánchez-Pérez et al., 2008), and larger seeds (Velasco et al., 2016), although the underlying genetic bases remain unknown, in particular for the two last traits.

In this study, we used population genomics to reconstruct the domestication and diffusion history of almond trees and identify the genes selected during domestication. We analyzed an extensive collection of cultivated almonds (*P. dulcis*) from Europe, North America, and Central Asia, as well as four wild almond relatives (*P. fenzliana*, *P. orientalis*, and *P. turcomanica* from the Caucasus and *P. spinosissima* from Central Asia). We also included back-to-wild almond tree individuals from New Zealand to investigate their recent diffusion history there. Using synonymous SNPs, we aimed to (1) characterize population structure, differentiation, and genetic diversity across wild and cultivated almond trees, (2) infer their divergence and demographic histories, and (3) identify genomic signatures of recent positive selection associated with domestication. Our results reveal high genetic diversity in cultivated almond trees, consistent with a weak domestication bottleneck and continued outcrossing. We also identify Central Asia as a hotspot for early domestication, and diversification from there towards the West in Europe and North America through massive crop-wild gene flow, in particular with the Oriental wild almond *P. orientalis*. We also detected region-specific selection signatures, but particularly in genes linked to the biosynthesis of the cyanogenic glycoside amygdalin and environmental adaptation.

## Materials and methods

### Samples, DNA extraction, and sequencing

For this study (Table S1), young leaves were collected from 30 wild almond trees: 13 *P. fenzliana* individuals native to the Caucasus and seven *P. spinosissima* individuals from Central Asia, all planted in a conservation orchard at INRAE, Bordeaux, France, and six *P. orientalis* individuals and four *P. turcomanica* individuals from East Anatolia, collected *in situ* from mother plants. Additionally, 34 *P. dulcis* cultivars were sampled from core collections in INRAE, Bordeaux and Avignon, and IRTA Constanti in Spain, including cultivars from Central Asia (*N*=7), the Caucasus (*N*=1), France (*N*=2), former USSR (*N*=1), Spain (*N*=16), and North America (*N*=7). Leaves of 18 *P. dulcis,* which naturally grew near Clyde in Central Otago and are supposed to be back-to-wild and invasive, were collected in New Zealand. Total genomic DNA was extracted using a standard rapid DNA isolation protocol (Doyle & Doyle, 1987). Sequencing libraries were prepared with a Nextera XT DNA Library Preparation Kit, and paired-end 150-bp reads (targeting 30× coverage) were generated using Illumina platforms (NextSeq 2000 and NovaSeq 6000).

This newly sequenced dataset was complemented by the previously published genome sequences of 14 *P. dulcis* from Central Asia (*N*=12) and North America (*N*=2) as paired-end ∼125-bp reads (about 60× coverage) (Yu et al., 2018), along with 11 additional *P. dulcis* accessions (nine from Europe and two from North America, Duval et al., 2023). In total, 109 almond samples were studied, comprising 82 newly sequenced and 25 previously published samples (Duval et al., 2023; Yu et al., 2018, Table S1). As an outgroup for specific analyses, five genotypes of *Prunus mira* Koehne sequenced in short reads at high coverage, the Tibetan wild peach (Cao et al., 2022), from a panmictic population in the Tibetan region (Table S1). The total dataset included 112 genotypes (Table S1).

### Read mapping and SNP calling

Analyses were performed using the *NYU IT High Performance Computing resources* platform (Figure S1). Read quality was assessed using FastQC v0.11.7 (Andrews et al., 2010). Reads were trimmed to 122–132 bp; reads less than 70 bp in length were removed using the fastp pre-processing tool v0.19.4. Individual reads were aligned to the almond reference genome, *P. dulcis* Texas genome v2.0 (Alioto et al., 2020), using bwa-mem2 v2.0 (Li, 2013) with the parameter: ‘mem –t 4’. SAMtools v1.9 (Li, 2011) and the ‘MarkDuplicates’ module from Genome Analysis Toolkit (GATK, v4.1.7.0) (https://software.broadinstitute.org/gatk/) (McKenna et al., 2010) were used to sort the mapped reads and remove redundant reads (i.e., PCR or optical duplicates). SNPs were then called using the ‘HaplotypeCaller’ module from GATK, setting ploidy to 2. The GATK modules ‘CombineGVCFs’ and ‘GenotypeGVCFs’ were used to combine the output and obtain one single variant call format (VCF hereafter) file for each of the eight chromosomes of the almond genome.

### SNP filtering

SNPs with missing rate > 10% and depth of coverage < 5 or >100 (i.e., sites with missing information) were removed with BCFtools v1.14 (Danecek et al., 2021) and the two GATK modules ‘VariantFiltration’ and ‘SelectVariants’. Insertion-deletion (InDel) polymorphisms were also removed. The cut-off for missing SNPs was determined by plotting the distribution of missing rate per sample in R version 4.1.2 (R Core Team, 2021) using the R packages ‘ggplot2’ (Wickham, 2016), ‘dplyr’ (Wickham et al., 2014), and ‘cowplot’ (Wilke, 2015). SNPs that became non-informative after the removal of those SNPs with a high missing rate, or too low or too high coverage, or with > 20% missing data, or with a minor allele frequency (MAF) < 0.05, were also removed using BCFtools. PLINK v2.0 (Chang et al., 2015) (https://www.cog-genomics.org/plink/2.0/) was used to detect duplicates, or closely genetically related individuals, setting the pairwise kinship-based inference for KING-robust kinship > 0.354. Highly genetically related individuals were identified with BCFtools, and all but one from each group of related samples were removed to avoid bias in population structure analyses.

For population structure and diversity analyses, non-synonymous and linked SNPs were further removed to avoid bias associated with confounding effects between demography and selection (Tiffin & Ross-Ibarra, 2014). First, SnpEff v5.1 (Cingolani et al., 2012) was used to annotate SNPs, and non-synonymous SNPs were removed using BCFtools v1.9 and SAMtools v1.9 (Danecek et al., 2021). Second, the remaining SNPs in linkage disequilibrium (LD) were removed. LD decay was estimated with PopLDdecay (Zhang et al., 2019) using the squared correlation coefficient (*r*²) between pairs of SNPs. The decayed physical distance between SNPs was estimated as the distance with half the maximum *r*² value. The decay of *r*² was plotted in R v4.1.2 (R Core Team, 2021) to determine a physical distance threshold below which SNPs were considered to be in linkage disequilibrium. Based on this threshold, VCFtools (Danecek et al., 2011) was used to thin the dataset by retaining a single SNP per LD block—specifically, the first SNP encountered within each interval exceeding the LD decay threshold.

### Population structure inferences

*The fast*STRUCTURE software v1.0 (Raj et al., 2014) was used to infer the population structure between cultivated and wild almond individuals, as well as among the cultivated almond individuals only, with the unlinked and synonymous SNP dataset defined above. This approach is based on the Hardy-Weinberg model and uses a Bayesian framework to infer panmictic groups for a predefined number of genetic clusters (*K*). The number of populations, *K*, was varied from two to ten, with 20 repetitions per *K* value. The program CLUMPAK (Kopelman et al., 2015) generated the consensus solution for each *K* value from the Q-matrices obtained by *fast*STRUCTURE. Results were annotated as major or minor clusters based on whether the population structure was detected by a majority or a minority of repetitions over the 20 total runs of *fast*STRUCTURE. Because *fast*STRUCTURE’s accuracy can be reduced under uneven sampling (Puechmaille, 2016), we evaluated this potential bias in our dataset. We used StructureSelector (Li & Liu, 2018), a web-based tool, to select the optimal K value more reliably. Population structure bar plots were generated using the R package ‘pophelper’ v2.3.1 (Francis, 2017). The distribution of the membership coefficient for the selected *K* value was plotted using the R package ‘ggplot2’ (Wickham, 2016). The cut-off value to assign each individual to a genetic cluster, as inferred by *fast*STRUCTURE, was determined based on this plot. Then, each individual was assigned to a cluster or considered admixed if the value was below the assignment threshold.

Two methods, without an *a priori* hypothesis, were used to investigate the genetic variation and differentiation among the genetic groups detected by *fast*STRUCTURE. First, a principal component analysis (PCA) was used to infer the genetic variation between individuals. The first three principal components were plotted with the R package ‘ggplot2’ (Wickham, 2016). Second, to estimate relatedness between individuals, a neighbor-net based on Nei’s distance between individuals (Nei, 1978) was built using PLINK v2.0 (Chang et al., 2015) and visualized using SplitsTree v4 (Huson & Bryant, 2006). For the PCA and neighbor-net, individuals assigned to a cluster were colored according to their genetic cluster. By contrast, admixed individuals (i.e., individuals with a membership coefficient below the assignment threshold) were colored grey.

### Genetic relationships, genetic diversity, and differentiation

For the subsequent analysis, admixed individuals and New Zealand individuals were removed from the dataset, as their samples did not represent a well-defined population (i.e., including only admixed individuals, see results below). Pairwise population differentiation (i.e., groups of individuals detected by *fast*STRUCTURE, excluding admixed individuals) was assessed using Weir and Cockerham’s *F_ST_* estimator, implemented in the Genepop software (version 1.2.2, Rousset, 2017). A global differentiation test across the 13,167 unlinked and synonymous SNPs was also conducted for each pair of populations using Fisher’s test for combining p-values. This approach sums the logarithms of individual locus-specific p-values into a *χ^2^* statistic with 2*k degrees of freedom (where k is the number of loci), allowing an overall significance test of genetic differentiation. The resulting *χ^2^* values and degrees of freedom were extracted from the Genepop output, and exact p-values were using the pchisq() function in the stats (version 3.6.2) R package (R Core Team, 2021). These p-values complement the *F_ST_* values by quantifying the statistical support for differentiation across loci. Heterozygosity (*H*_o_, observed heterozygosity; and *H*_e_, expected heterozygosity), and the inbreeding coefficient for each population were estimated with Stacks v2.61 (Catchen et al., 2013). Genetic diversity (π) and genetic distance (*D_XY_*) between populations were estimated using Pixy (Korunes & Samuk, 2021) with the “all-sites” VCF file. Genome-wide π and *D_XY_* were calculated using all sites (332,335,748 variant and invariant sites) in 10-Kb non-overlapping windows of high mappability. The nucleotide diversity (π) values were compared between populations using the Mann-Whitney-Wilcoxon U test (Mann & Whitney, 1947; Wilcoxon, 1945).

To investigate the relationship among the populations, a species tree was generated using the Singular Value Decomposition Scores for Species Quartets, SVDQuartets (Chifman & Kubatko, 2014). The Tibetan wild peach (*P. mira*, Guanghetao accessions) was used as an outgroup for the tree (*N* = 5) (Cao et al., 2022). The same method described above was used to prepare a new SNP dataset with these five wild peach genomes. Synonymous and unlinked SNPs were used for this analysis. SNPs were filtered again, using BCFtools, to remove SNPs with more than 20% missing data, SNPs with MAF < 0.05, or SNPs that became non-informative after the removal of other SNPs. Then, the resulting VCF file was converted to a Nexus file using the Ruby script provided on the SVDQuartets GitHub page (https://github.com/ForBioPhylogenomics/tutorials/tree/main/species_tree_inference_with_sn <u>p_data</u>). The module PAUP4 (paup-v4a166) was used to set the outgroup as *P. mira* genomes and launch SVDQuartets. The FigTree v1.4.4 tool (Rambaut, 2009) was used to visualize the tree obtained with SVDQuartets.

### Effective population size variation and divergence time between populations

A multiple sequentially Markovian coalescent (MSMC2) model was employed to investigate the demographic history of almond populations by estimating the variation in effective population size (*Ne*) over time and the timing of separation between the two populations (Schiffels & Wang, 2020). MSMC is an algorithm and program that analyzes multiple-phased genome sequences simultaneously (separated into haplotypes, i.e., maternal and paternal haploid chromosomes) to fit a demographic model to the data. It is an extension of pairwise sequentially Markovian coalescent (PSMC) to multiple individuals.

Haplotypes were randomly selected for each population: three genomes were randomly picked, three times in each population, to obtain three replicate analyses per population. The VCF files were phased using the software package Beagle 5.1 (version 18May20.d20) (Browning et al., 2018) with default parameters (no genetic map, window = 1, and overlap = 0.1). BCFtools v1.9 (Danecek et al., 2021) was used to extract individual information from the VCF file for the genomes previously selected, and to remove sites with ‘F_MISSING > 0 || AC == 0’. The sequence mappability of the *P. dulcis* Texas genome v2.0 was computed with GenMap 1.3.0 (https://github.com/cpockrandt/genmap) (Pockrandt et al., 2020) using the parameters ‘-K 140 –E 0 –T 40’, which assesses the uniqueness of all possible 140-mers in the genome, allowing no mismatches. A custom Perl script (findmask.pl) was used to obtain the sorted and merged.bed file with the low-mappability regions of the genome masked, using a window of 20 kb and a step size of 10 kb. The complement bed files for each masked chromosome were obtained using the ‘complementBed’ module of Bedtools v2.25.0 (Quinlan & Hall, 2010). Finally, the MSMC program ‘generate_multihetsep.py’, which merges the VCF files and the mask complement files, was used to generate the input file for MSMC2.

The time segment patterning option of MSMC2 (msmc2-v2.1.2) was set to ‘-p 1*2+16*1+1*2’. First, MSMC2 was run to estimate the coalescence rate within each almond population, thereby inferring *N*_e_ variation. Then, it was run to estimate the coalescence rate across populations, thereby inferring the population divergence history. The results were plotted in R version 4.1.2 (R Core Team, 2021) using the R packages ‘ggplot2’ (Wickham, 2016), ‘scales’ (Wickham et al., 2011), ‘grid’ (Murrell & Wen, 2014), and ‘cowplot’ (Wilke, 2015). The generation time was set to 10 years, and the substitution rate was set to 10^−8^ nucleotides per site per year (Velasco et al., 2016). Some geological time markers were added to the plot: the last and penultimate glacial periods, the Pleistocene epoch, and the Holocene epoch.

### Gene flow occurrence between crop and wild populations

Setting *P. mira* as the outgroup, an ABBA-BABA test was performed between all possible triplets among individuals of *P. dulcis* from Central Asia, Europe (clusters I and II), and North America, *P. fenzliana*, *P. orientalis*, and *P. spinosissima* populations (i.e., groups of individuals detected by *fast*STRUCTURE, excluding admixed individuals), using the Dtrios program within Dsuite v0.5r48 (Malinsky et al., 2021). The input VCF file was the same as for the SpeciesTree: *fast*Structure input (unlinked synonymous SNPs), excluding admixed individuals and those from New Zealand, and including *P. mira* individuals (*N* = 5).

### Detection of positive selection in the genomes

To detect putative signatures of positive selection (i.e., selective sweeps), we focused on the almond populations identified through *fast*STRUCTURE (see Results). A selective sweep describes the rapid increase in frequency of a beneficial allele, often resulting in elevated LD and reduced nucleotide diversity in the surrounding genomic region. To minimize confounding effects from admixture, admixed individuals (assignment probability < 0.85, see results) were excluded using BCFtools v1.9 (Danecek et al., 2021).

Multiple complementary approaches were implemented to identify genomic regions under selection. First, RAiSD (Alachiotis & Pavlidis, 2018) was used to calculate the composite statistic *μ*, which integrates variation in nucleotide diversity, LD, and the site frequency spectrum to identify candidate sweep regions. Second, OmegaPlus v2.3 (Alachiotis et al., 2012) was applied to estimate the *ω* statistic, a model-free measure based on LD decay around candidate sites, which allows for the detection of both hard and soft sweeps. In addition, population-specific analyses of selection were performed on *P. dulcis* populations using the population branch statistic (PBS), which quantifies changes in lineage-specific allele frequency by comparing a focal population to two reference populations (Yi et al., 2010). The PBS enables the detection of selection signatures unique to the focal population while taking into account past demographic history.

To generate low-mappability masks prior selection scans, we employed SNPable to remove unreliable genomic positions. This method simulates 35 bp k-mers from the reference genome and maps them back using BWA (Li, 2013), retaining only uniquely mappable positions with at least 99% identity. The resulting high-mappability regions were extracted as a BED file and used to filter out poorly mappable regions before downstream analyses. Compared to genmask, which was used in our demographic analyses (MSMC2) to exclude repetitive or low-complexity genomic regions, SNPable provides a more fine-scale and read-mapping-aware estimation of mappability. Notably, genmask yielded considerably more restricted mappable regions, which could lead to excessive masking in selection scans. Therefore, SNPable was chosen here to achieve a better balance between mapping reliability and genome coverage.

After removing SNPs located in regions with low mappability, we retained 6,063,769 of 8,132,54 SNPs initially detected, for sweep detection using OmegaPlus and RAiSD estimates, and 744,241 SNPs for PBS estimates (as we only kept common SNPs to the populations compared, see results). Genome-wide significance thresholds were set to the 99th percentiles. Selection scan plots were generated using the package ‘qqman’ in R (Turner, 2014). Functional gene annotations were retrieved from eggNOG-mapper (Cantalapiedra et al., 2021) annotations (v5.0) of the *Prunus dulcis* reference genome (Texas cultivar), which included GO terms assigned to each predicted gene. Individual and intersecting gene annotations were visualized with the R packages ‘UpSetR’ (Conway et al., 2017) and ‘ComplexUpset’ (Krassowski, 2020).

To investigate whether genes under positive selection were enriched for particular biological functions, we performed Gene Ontology (GO) enrichment analyses using the clusterProfiler R package (Yu et al., 2012). For each almond population (PdCA, PdE1, PdE2, PdNA, Po, Pf and Ps), we identified genes containing the top 1% outlier of the selection statistics, detected using at least two methods across RAiSD, OmegaPlus and PBS. GO enrichment was conducted using the enricher() function, testing the selected genes against the full set of annotated genes as background. Only GO terms with a q-value (FDR-adjusted p-value) below 0.05 were retained.

## Results

### SNP filtering and final dataset

Following mapping of sequencing reads onto the *P. dulcis* Texas Genome v2.0 reference almond genome, SNP calling, and quality control, the resulting dataset consisted of 8,132,541 high-quality SNPs for 96 individuals (excluding 11 closely genetically related individuals or low-quality samples, Table S1), representing ∼1% of the almond genome size, with a mean SNP density of one SNP every 111 bp. We included 59 cultivated almond individuals (*P. dulcis*), nine back-to-wild invasive New Zealanders, and 28 wild almond individuals, with six *P. orientalis*, four *P. turcomanica*, 12 *P. fenzliana*, and six *P. spinosissima* (Figure 1A).

**Figure 1.**
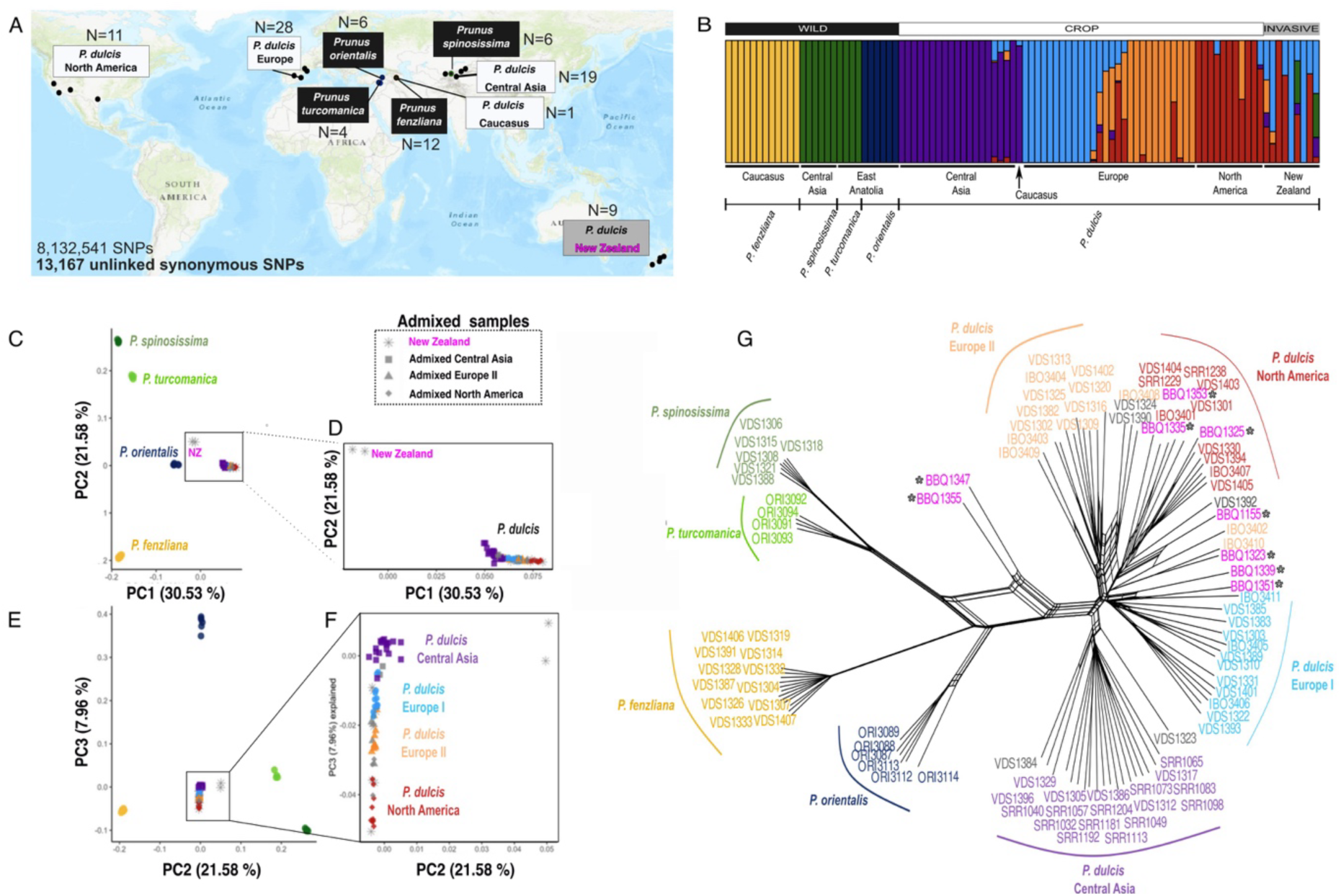
Population genetic structure and relationships among wild and cultivated almond. (A) Geographic origins of the sampled accessions: cultivated Prunus dulcis [N = 59 from Central Asia (N = 19), the Caucasus (N = 1), Europe (N = 28) and North America (N = 11)], back-to-wild supposed invasive P. dulcis from New Zealand (N = 9), Prunus fenzliana (N = 12), Prunus orientalis (N = 6), Prunus turcomanica (N=4), and Prunus spinosissima (N = 6). Species names are color-coded to indicate crop (white), wild (black), or back-to-wild (grey) status (i.e., New Zealand samples in pink). (B) Population structure inferred with fastSTRUCTURE at K = 8 (N = 96 individuals after removing closely genetically related individuals; 13,167 unlinked synonymous SNPs), with samples grouped by species and geographical origin (Caucasus, Central Asia, East Anatolia, Europe, North America, and New Zealand). Each vertical bar represents one individual, and color segments represent ancestry proportions from each inferred genetic cluster. Although eight clusters were specified, only seven are visually distinguishable. (C–F) Principal component analyses (PCA) based on 13,167 genome-wide unlinked and synonymous SNPs. A total variance of 60.1% is explained by PC1 (30.5%), PC2 (21.6%), and PC3 (8.0%). (C) PC1 and PC2 plot, (E) PC2 and PC3 plot for all samples (N = 96). (D) and (F) Zoomed-in views of cultivated almonds and admixed individuals (N = 61), highlighting finer-scale population structure. (G) Neighbor-net based on pairwise genetic distances among all accessions, confirming the genetic differentiation between wild species and cultivated almonds, and revealing relationships among P. dulcis populations inferred with fastSTRUCTURE. Colors correspond to populations as shown in (B) and admixed individuals are colored in grey; all New Zealanders were admixed and indicated by a grey star* and in pink in the plots.

To ensure a robust analysis of population structure, genetic diversity, and differentiation, we applied additional filtering steps to minimize the confounding effects of natural selection and LD. We retained 131,001 synonymous SNPs after excluding non-synonymous SNPs. Based on an estimated LD distance of ∼8 Kb (corresponding to a drop in r² from ∼0.23 to ∼0.12; Figure S2), we thinned the dataset by retaining only one SNP within each 8 Kb window. Specifically, the genome was scanned in non-overlapping 8 Kb intervals, and one representative SNP per window (typically the first encountered) was retained, resulting in a final dataset of 13,167 unlinked synonymous SNPs (Figure 1A), which served as the basis for population structure and diversity analyses.

### Three wild and four cultivated populations spread across Eurasia and North America

*fast*STRUCTURE identified seven main genetic clusters among wild and cultivated almond individuals spanning Eurasia and North America (Figure 1B). The cross-validation error diminished steadily until reaching *K* = 10, and StructureSelector indicated that the optimal number of clusters lay between *K* = 6 and *K* = 9 (Figure S3). For *K* = 8, we detected multiple clustering modes using CLUMPAK, with the third most frequent configuration (Minor cluster 2) providing the best match to known biological and geographical distributions (Figure S4). Despite being computed with *K* = 8, this mode revealed seven distinguishable clusters, suggesting that the eighth component likely resulted from model overfitting. For *K* = 8 (Minor cluster 2), cultivated *P. dulcis* individuals were divided into four major regional groups: Central Asia, North America, Europe I (primarily Spanish cultivars), and Europe II (Spanish, French, and Italian cultivars). The wild species formed three distinct clusters: *P. fenzliana*, *P. orientalis*, and a group combining *P. spinosissima* and *P. turcomanica* (green). Accessions from New Zealand (*P. dulcis*) were highly admixed, likely drawing ancestry from multiple sources. These individuals showed significant ancestry from the European (Europe I and II) and North American cultivated groups, and to a lesser extent from *P. spinosissima*. Only three New Zealand individuals showed unambiguous assignment to a single cluster: one grouped with North American *P. dulcis* and two with Europe I. A *fast*STRUCTURE analysis restricted to the cultivated subset (*N* = 68 individuals, using the 13,167 unlinked synonymous SNPs, Figure S5) produced clustering results consistent with those from the full dataset (Figure S4), supporting the robustness of the seven-group structure. We retained this clustering pattern for *K* = 8 (Minor cluster 2 configuration), which was inferred using the whole dataset, for all downstream analyses.

We assigned each individual to a cluster using a membership coefficient threshold of 0.85 (Figure S6, Table S2), resulting in 80 individuals confidently assigned to one of the seven populations, whereas 16 individuals, including eight genotypes from New Zealand (Tables S1 and S2), were classified as admixed.

The PCA and neighbor-net based on 13,167 unlinked synonymous SNPs confirmed (1) the seven populations identified by *fast*STRUCTURE, but further revealed a genetic subdivision, distinguishing *P. spinosissima* and *P. turcomanica,* see below); (2) the subdivision of cultivated almond individuals into four populations corresponding to their geographical origins, with a notable split between southern Spanish (European II) and other European accessions (European I); and (3) that New Zealand back-to-wild almond individuals did not form a distinct cluster but were admixed with Europe I and North American *P. dulcis* individuals. A PCA distinguished the *P. dulcis*, *P. fenzliana*, *P. orientalis*, *P. turcomanica*, and *P. spinosissima* along the first two principal components 1 and 2 (30.5% and 21.6% of explained variance, respectively; Figure 1C and D). Cultivated almond individuals (*P. dulcis* from Europe, North America, Central Asia, and New Zealand), assigned to four populations with *fast*STRUCTURE (admixed in grey), clustered apart from those of wild species along PC1 and PC3 (Figures 1E and F). New Zealand almond individuals overlapped with those of European and North American cultivars in the PCA. The neighbor-net (Figure 1G) mirrored these results, recovering distinct clusters for the three wild species individuals and a cultivated almond clade split into four geographically defined populations. Central Asian *P. dulcis* individuals were genetically different from those of the other cultivated groups, whereas New Zealand almond individuals again clustered with European and North American *P. dulcis* individuals, reflecting their admixed origin. The neighbor-joining tree also revealed a clear split between *P. turcomanica* and *P. spinosissima*, supporting their treatment as distinct populations. However, due to the limited sample size for *P. turcomanica* (only four individuals), this group was excluded from downstream analyses. Consequently, we focused on seven populations: four cultivated *P. dulcis* populations (Europe I, Europe II, North America, and Central Asia) and three wild almond populations (*P. orientalis*, *P. spinosissima*, and *P. fenzliana*) (Tables 1 and 2).

**Table 1.**
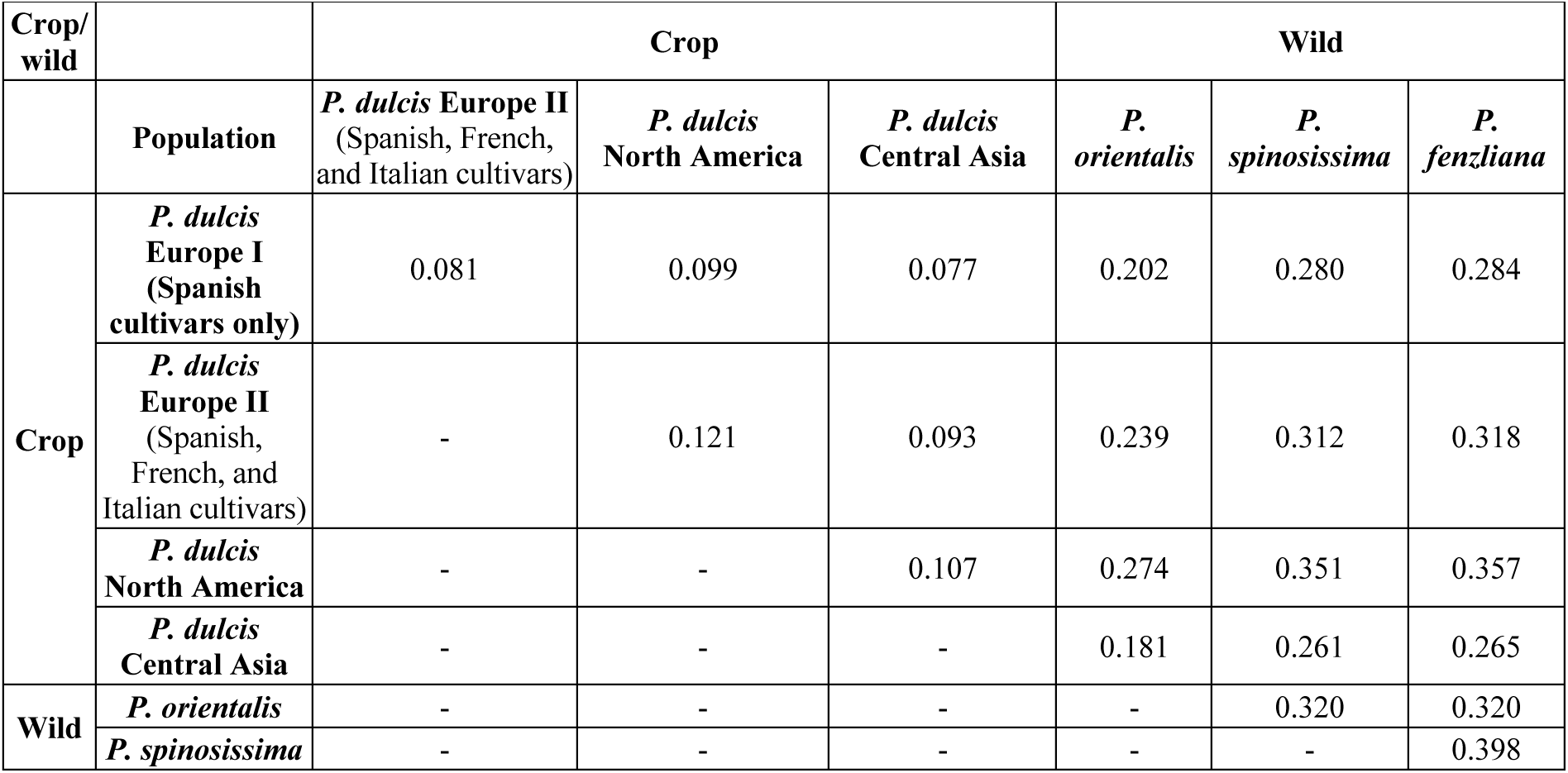
Genetic differentiation estimates (*F*_ST_) among wild and cultivated almond populations. Populations were identified using *fast*STRUCTURE with *K* = 8 (*N* = 77; admixed individuals and New Zealand almond individuals were excluded, as well as *P. turcomanica*; 13,167 unlinked synonymous SNPs). All *F*_ST_ values were significant (*P* < 0.001).

**Table 2.** Genetic diversity estimates for wild and cultivated almond populations. Populations were identified by *fast*STRUCTURE with *K* = 8 (*N* = 77; admixed individuals and New Zealand almond individuals were excluded; 13,167 unlinked synonymous SNPs). Diversity metrics were computed using Stacks v2.68 and Pixy v2.0.0.

| Almond population | N | SNPs | Private alleles | $H_o$ | $H_e$ | $\pi$ | $F_{IS}$ |
| --- | --- | --- | --- | --- | --- | --- | --- |
| *P. fenzliana* | 10 | 3,732 | 1,361 | 0.111 | 0.094 | 1.48e-5 | -0.025 |
| *P. spinosissima* | 6 | 2,760 | 664 | 0.083 | 0.071 | 1.07e-5 | -0.009 |
| *P. orientalis* | 10 | 6,154 | 270 | 0.0998 | 0.160 | 2.32e-5 | 0.148 |
| Central Asia *P. dulcis* | 20 | 7,621 | 276 | 0.175 | 0.165 | 2.39e-5 | -0.007 |
| North America *P. dulcis* | 6 | 4,872 | 49 | 0.204 | 0.139 | 2.36e-5 | -0.093 |
| Europe I *P. dulcis* | 8 | 6,749 | 78 | 0.199 | 0.168 | 2.55e-5 | -0.041 |
| Europe II *P. dulcis* | 7 | 5,733 | 95 | 0.188 | 0.153 | 2.51e-5 | -0.048 |

$N$ , number of individuals assigned to a population (group of individuals assigned with a membership coefficient $> 0.85$ to a given cluster); SNPs, number of SNPs retained per population; $H_o$ , observed heterozygosity; $H_e$ , expected heterozygosity; $\pi$ , nucleotide diversity; $F_{IS}$ , inbreeding coefficient.

### An older separated Central Asian cultivated population, a wild ancestry from *P. orientalis,* and crop-wild gene flow driving Western almond diversification

Genetic differentiation estimates revealed that cultivated almond populations were weakly differentiated from one another (Table 1). By contrast, divergence between wild and cultivated almond populations was substantially higher (*F*_ST_ = 0.26–0.43), with the greatest differentiation observed between the *P. fenzliana* and *P. spinosissima* populations (*F*_ST_ = 0.50). SVDquartet tree further clarified these relationships, confirming *P. orientalis* as the genetically closest wild relative to *P. dulcis*, whereas *P. fenzliana* and *P. spinosissima* individuals appeared more distantly related (Figure 2A). Among cultivated almond populations, the Central Asian *P. dulcis* population formed a distinct clade, followed by North American almond population. In contrast, European almond populations (Europe I and II) clustered into a more recently diverged group, albeit with low bootstrap support, suggesting unresolved relationships likely shaped by recent divergence or gene flow.

**Figure 2.**
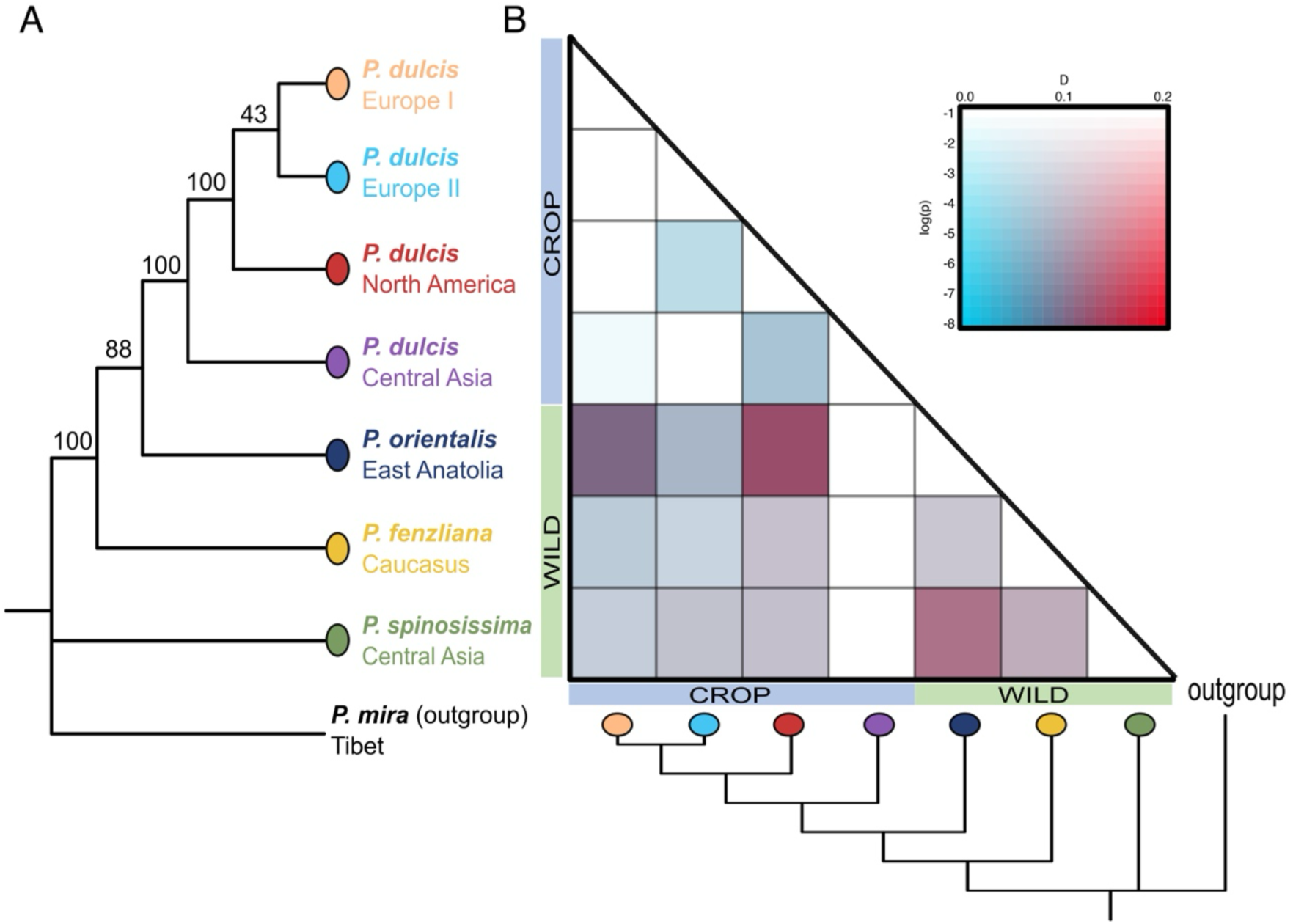
Phylogenetic relationships and gene flow among cultivated almonds and wild relative populations. **(A)** Phylogenetic tree based on putatively synonymous and unlinked SNPs, illustrating relationships among cultivated *P. dulcis* populations (Europe I, Europe II, North America, Central Asia) and wild species (*P. orientalis*, *P. fenzliana*, *P. spinosissima*), with *P. mira* used as an outgroup. Bootstrap support values are indicated at each node. **(B)** Gene flow test (*D*-min estimates) among almond populations, using *P. mira* as the outgroup. Blue indicates lower occurrences of gene flow, red indicates higher occurrences, with darker shades reflecting higher statistical significance. Population labels correspond to those detected with *fast*STRUCTURE at *K*=8 (i.e., groups of individuals assigned with a membership coefficient >0.85).

ABBA-BABA analysis revealed the occurrence of pervasive wild-crop gene flow (Figure 2B), particularly between *P. orientalis* with the European and North American cultivated almonds. By contrast, gene flow between wild species and the Central Asian almonds was minimal, suggesting a more isolated domesticated gene pool in this region. These patterns stand in contrast to the relatively low genetic differentiation among cultivated almonds, underscoring the asymmetric contributions of wild gene pools to domesticated almonds across areas. Together, these findings suggest that whereas almond individuals from the Central Asian group may reflect an early domestication lineage with limited external introgression, European and North American almond groups have been substantially shaped by admixture with local wild relatives, most likely during their westward and overseas diffusion.

### Genetic diversities of cultivated almond populations indicate various domestication and breeding histories, and demographic histories shaped by past climate changes

Genome-wide diversity analyses provided insights into the global and regional domestication history of almonds. Cultivated almond individuals (*P. dulcis*) retained significantly more genetic diversity than their wild relatives (Table 2, *P*<0.001; Table S3), particularly compared to *P. fenzliana* and *P. spinosissima*, consistent with a weak domestication bottleneck characteristic of perennial fruit crops. Within cultivated populations, the lowest diversity was observed in the North American population, also characterized by the highest observed LD pattern (Figure S7) and a lower number of private alleles, indicating a recent origin consistent with a transcontinental introduction (Decroocq et al., 2025). The highest diversity was observed in the European cultivated populations, which also exhibited an intermediate number of private alleles is consistent with recent westward dispersal. *F_IS_* values were all negative in all cultivated almond populations, consistent with predominantly outcrossing mating system which favours gene flow and lack of inbreeding. The Central Asian *P. dulcis* preserved the highest number of private alleles, indicative of a possible early domestication center.

In the wild populations, *P. orientalis* displayed contrasting patterns compared to the other wild species. The Oriental wild almond showed the lowest number of private alleles but the highest genetic diversity and a positive inbreeding coefficient (*F_IS_* = 0.148), suggesting the presence of cryptic substructure, but lack of a sufficient number of samples could not help to dig further into it. Among wild almond individuals, *P. fenzliana* possessed the greatest number of private alleles (1,361 sites), followed by *P. spinosissima* (664 sites), validating their highest genetic differentiation (Table 1 and Figure 2).

MSMC2 demographic inferences (Figure 3) indicate that both wild and cultivated almond populations underwent a historical increase in *N*e during the Mid-Pleistocene, followed by a marked contraction around the Last Glacial Maximum (LGM) and a subsequent post-glacial expansion. Despite its low present-day nucleotide diversity, *P. fenzliana* exhibited signals of a large ancestral *N*e in MSMC2 analyses. This apparent discrepancy suggests that its reduced genetic diversity (Table 2) likely reflects a recent population contraction or sampling from genetically isolated populations, rather than a long-term small *N*e. The exceptionally high number of private alleles in *P. fenzliana* (n = 1,361; Table 1) further supports a history of prolonged genetic isolation and deep divergence, reinforcing its uniqueness as a wild gene pool. For cultivated almonds, MSMC2 trajectories suggest somewhat lower historical *N*e values compared to wild relatives, with the lowest estimates observed in the North American population—consistent with its reduced genetic diversity and slower LD decay. However, these results should be interpreted cautiously: almond domestication, like that of other perennial fruit trees, is thought to be relatively recent (∼4,000 years ago, (Zohary & Spiegel-Roy, 1975), and MSMC2 is known to be less reliable at resolving demographic events in the last ∼1,000 years (Santiago et al., 2020). Nevertheless, the observed trends align with patterns of genetic diversity, supporting lower *Ne* in western populations than in Central Asia, a region of retained ancestral diversity.

**Figure 3.**
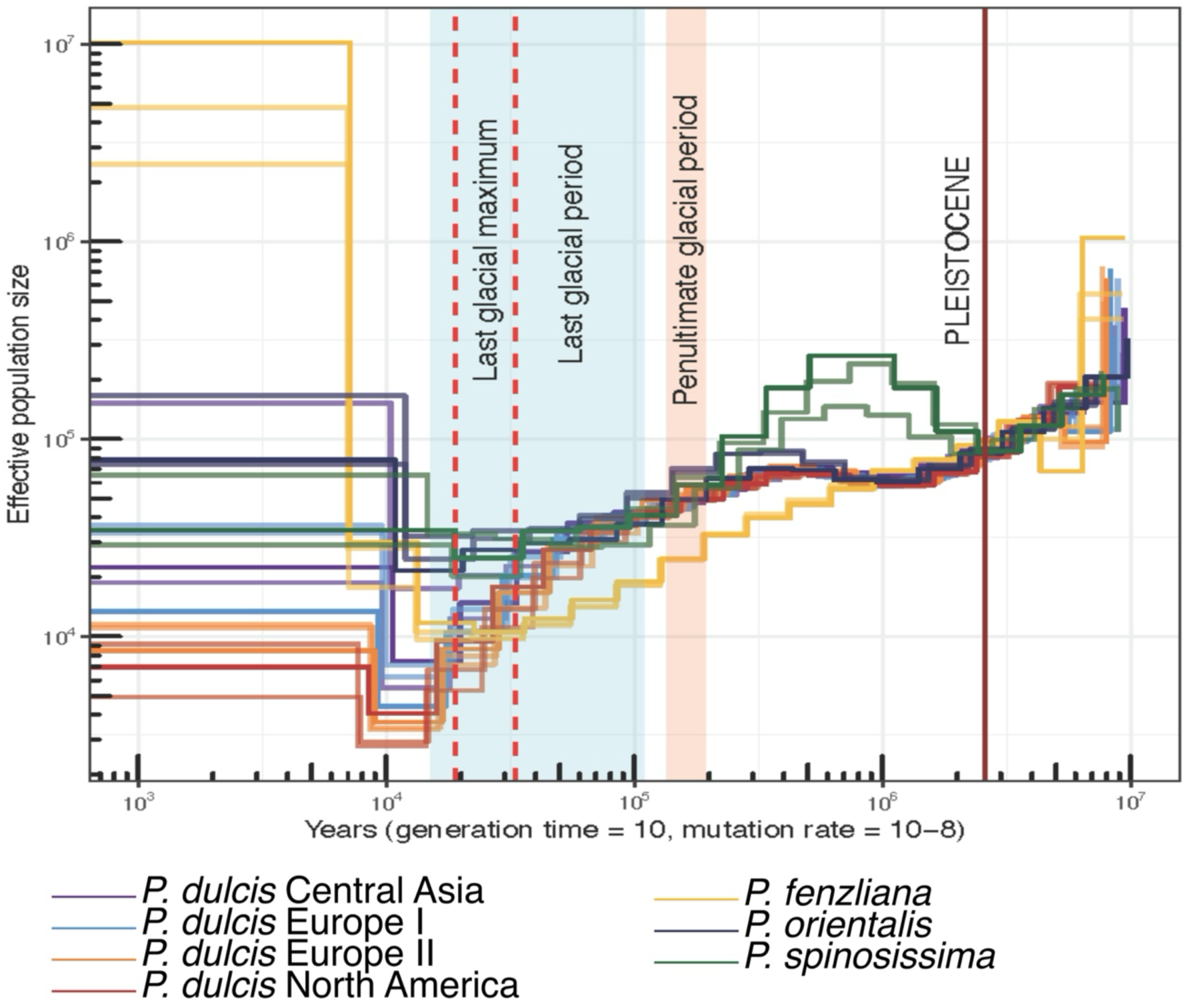
Historical effective population size (*N*_e_) inferred using MSMC2 for cultivated and wild relative almond populations. The *x*-axis represents time in years (assuming a generation time of 10 years and a mutation rate of 10^−8^). Major climatic periods are highlighted: the penultimate glacial period (pink) and the last glacial maximum (blue). Vertical red dashed lines mark the approximate beginning and end of the last glacial period. For these analyses, *N* = 77; admixed individuals and New Zealand almond individuals were removed; 13,167 unlinked synonymous SNPs. Due to the limited sample size for *P. turcomanica* (only four individuals), this group was excluded from analyses following population structure inferences (see results above).

### Candidate genes under selection and shared targets across cultivated almond populations

Signals of selective sweeps were distributed heterogeneously across the genome (Figures 4 and S8, Table S4). Chromosome 1, being the largest in the almond genome, contained the highest number of candidate genes under selection. In all cultivated almond populations, the selective sweep signatures were scattered across chromosomes without pronounced clustering. Distinct, population-specific selection patterns were also observed. For example, chromosome 7 showed strong selection signals in the North American population, while chromosome 1 was a key target in Central Asian almonds, reflecting regional differences in selective pressures during domestication.

**Figure 4.**
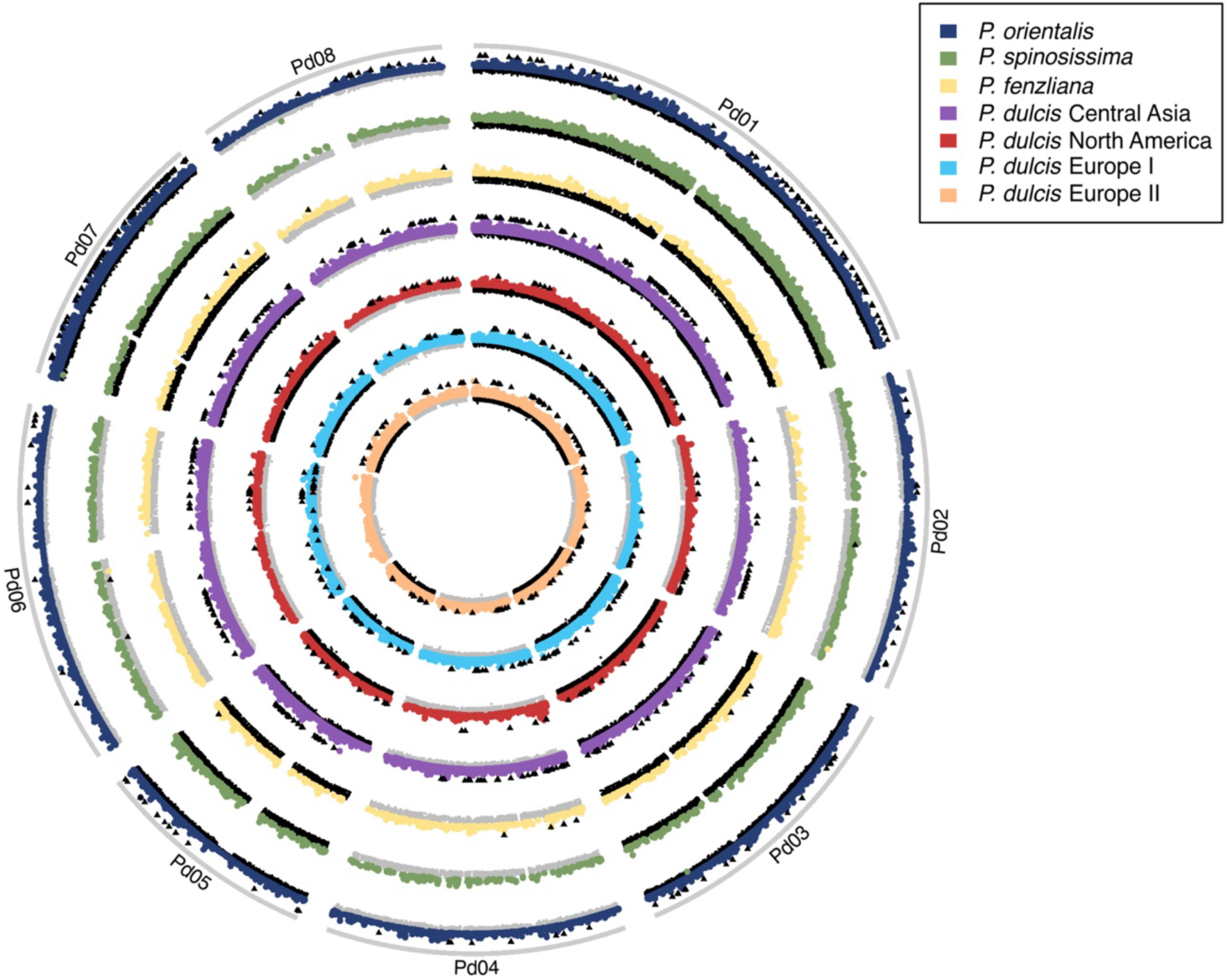
Genome-wide detection of selective sweeps in cultivated and wild almond populations. Selective sweep analysis was performed across the whole genomes of cultivated and wild almond populations using the OmegaPlus and RAiSD methods, with a set of 6,063,769 SNPs. Black triangles indicate population-specific genes showing selective sweep signatures using at leat two methods (numbers of genes per population: *PdCA*, 339; *PdE1*, 218; *PdE2*, 164; *PdNA*, 147; *Pf*, 19; *Po*, 224; *Ps*, 41).

Cross-referencing selective sweep regions in cultivated and wild populations with genome annotations identified between 1,245 and 4,565 candidate genes per population by at least one of the three sweep detection methods used (Figure S9). The lower number of genes detected in *P. spinosissima* and *P. fenzliana* was likely due to the exclusion of PBS for these wild populations. For the cultivated almond populations, we applied PBS to detect genes specifically selected during almond domestication, while explicitly accounting for the gene flow and divergent history inferred above (Figures 2 and 3). PBS uses a three-population framework, where a focal population is compared to two reference populations, allowing the identification of lineage-specific allele frequency changes that are not confounded by shared demographic events. We therefore used the populations inferred from *fast*STRUCTURE: a focal cultivated population (either European I, European II, or North American), and *P. dulcis* from Central Asia and *P. orientalis* as the two reference populations (Figure S10). This setup was justified by our earlier analyses (Figure 2; Tables 1 and 2), which showed that *P. orientalis* was the genetically closest wild relative to cultivated almonds, while Central Asian *P. dulcis* was the most genetically diverse and divergent cultivated group. To increase confidence in selection signals, we focused on genes detected by at least two complementary approaches (Figure S11).

We first examined the few shared signals of selection among cultivated almond populations detected by at least two complementary approaches (Table S4). Notably, the two European *P. dulcis* populations shared 13 selective sweep regions (Figure S11). In contrast, only 10, 4, and 2 genes with sweep signals overlapped between the North American population and the Central Asian, European I, and European II almonds, respectively. This limited overlap indicates that the selective signals detected in the North American population primarily reflect its unique selection history, rather than domestication events shared across regions. In total, 122 (9.6%) of these genes showed signatures of selection in at least two almond populations (Figure S11, Table S4) and 64 (5.0%) were shared across at least two cultivated populations (Figure S11, Table S4), making them strong candidates for key domestication-related targets, involved in traits such as stress response, flowering time regulation, or kernel quality. Notable candidate genes included those involved in biological defense pathways. Genes encoding a glutamate receptor (Prudul26A013903) and a leucine-rich repeat (LRR) receptor-like serine-threonine protein (Prudul26A029457, Prudul26A028737)—two types of proteins involved in calcium signaling and plant immune responses in *Arabidopsis thaliana* (Lee et al., 2025). We also highlighted domestication-related traits such as aroma and kernel sweetness. A gene encoding a strictosidine synthase–like protein (Prudul26A026786), involved in the indole alkaloid biosynthesis pathway and aroma production in crops like lemon (*Citrus* sp.) (Gonzalez-Ibeas et al., 2021) and sweet potato (Qiao et al., 2023), also showed selection signatures across cultivated almond populations. Notably, in cultivated apples, calcium treatment increases the abundance of fatty acid metabolism–related enzymes, such as Strictosidine synthase–like 5 (STR5), which influences aroma volatile production (Xu et al., 2022).

To assess whether *P. orientalis*, the genetically closest wild relative of *P. dulcis*, contributed adaptive alleles to the cultivated almond genome, we compared selective sweeps detected in *P. orientalis* with those identified in each cultivated population. Overall, shared selective footprints were unevenly distributed across cultivated populations, suggesting differences in the extent of historical gene flow or retention of ancestral polymorphism. The Central Asian *P. dulcis* population (PdCA)—considered the likely cradle of almond domestication—shared the largest number of selective sweep genes with *P. orientalis* (30 genes). For instance, two genes encode proteins homologous to those previously shown to be involved in the amygdalin pathway: Prudul26A014980, which encodes a cytochrome P450 protein, and Prudul26A024658, which encodes a UDP-glycosyltransferase. Homologous proteins have been shown to play crucial roles in bitterness (Thodberg et al., 2018). Among these genes, we also found that several could play roles in seed yields, including Prudul26A015909 and Prudul26A021806, which encode a sulfate transporter. These genes could play a role in almond domestication, as they were described as determinants to enhance seed yield in pea (Bachelet et al., 2024). Finally, Prudul26A005915 encodes a sucrose-cleaving enzyme that provides UDP-glucose and fructose for various metabolic pathways. Given the well-documented role of sucrose as both a metabolic substrate and a key signaling molecule in flowering induction, selection acting on this gene may reflect adaptation in the regulation of flowering time. Although this gene lies outside the region defined by the previously reported flowering time QTL associated with the CPSCT042 marker (∼15.5 Mb on Pd07, Sánchez-Pérez et al., 2012), its potential functional relevance warrants further investigation. The European I population (PdE1) shared nine genes under selection with *P. orientalis,* whereas the European II population (PdE2) shared eight genes. In each of these European populations, we found, among them, genes encoding enzymes likely involved in the amygdalin pathway (Prudul26A026629 encoding a UDP-glycosyltransferase in PdE1 and Prudul26A032640 encoding a cytochrome P450 in PdE2). Finally, the North American almond population (PdNA) exhibited the fewest shared genes (5) with *P. orientalis*. Among them, Prudul26A031075 encoding a sugar transporter, could be linked to biotic and abiotic stress responses (Julius et al., 2017; Saddhe et al., 2021). Together, these results highlight a gradient of shared selection signals, with the strongest signals observed in Central Asian almonds (east) and the weakest in North America (west). Focusing on amygdalin pathway, we observed that, in addition to the previously mentioned genes showing selective sweep signatures, seven genes (Prudul26A030744, Prudul26A019284, Prudul26A020338, Prudul26A005388, Prudul26A015550, Prudul26A009250, Prudul26A022082 encoding cytochrome p450) exhibited strong selection signals in at least one of the cultivated populations, and for Prudul26A030744 and Prudul26A020338, in the *P. orientalis* (Table 3). However, none of the selective sweep signals in these genes are shared by all cultivated populations and none of them are overlapping with previously identified candidate genes (Alioto et al., 2020; Thodberg et al., 2018), suggesting a regionally variable genetic basis for the sweet kernel phenotype, consistent with previous suggestions (Alioto et al., 2020; Sánchez-Pérez et al., 2019). In addition, seven genes encoding tobacco mosaic virus (TMV) resistance proteins (e.g., Prudul26A000912, Prudul26A013852, Prudul26A015345, Prudul26A016154, Prudul26A022384, Prudul26A023571, Prudul26A024982) showed sweep signals in one to three cultivated populations, depending on the gene considered, and for two genes (Prudul26A000912 and Prudul26A013852), in *P. orientalis* (Table S4).

**Table 3.** Genes encoding enzymes involved in the amygdalin pathway, which affects almond kernel bitterness, showing positive selection signatures in wild and cultivated almond populations, detected using at least two methods.

| Gene ID | Description | Chromosome | Population | Method |
| --- | --- | --- | --- | --- |
| Prdul26A026629 | Belong to the cytochrome p450 family | Pd01 | *P. orientalis*, *P. dulcis* from Central Asia | RAiSD, OmegaPlus |
| Prdul26A032640 | Belong to the cytochrome p450 family | Pd07 | *P. orientalis*, *P. dulcis* from Europe II | RAiSD, PBS |
| Prdul26A014980 | Belong to the cytochrome p450 family | Pd06 | *P. orientalis*, *P. dulcis* from Central Asia | OmegaPlus, PBS, RAiSD |
| Prdul26A024658 | Belongs to the UDP-glycosyltransferase family | Pd01 | *P. orientalis*, *P. dulcis* from Central Asia | RAiSD, PBS |
| Prdul26A030744 | Cytochrome p450 | Pd01 | *P. orientalis*, *P. dulcis* from Europe I and II | OmegaPlus, PBS, RAiSD |
| Prdul26A020338 | Belong to the cytochrome p450 family | Pd01 | *P. orientalis*, *P. dulcis* from Central Asia, North America | RAiSD, PBS |
| Prdul26A005388 | Belong to the cytochrome p450 family | Pd01 | *P. dulcis* from Europe I and II | OmegaPlus, PBS, RAiSD |
| Prdul26A019284 | Belong to the cytochrome p450 family | Pd01 | *P. dulcis* from Central Asia | RAiSD, PBS |
| Prdul26A009250 | Cytochrome p450 | Pd02 | *P. dulcis* from Europe II | OmegaPlus, RAiSD |
| Prdul26A022082 | Belong to the cytochrome p450 family | Pd08 | *P. dulcis* from Europe II | RAiSD, PBS |

Most of the genes under selection detected in at least two methods were population-specific (Figures 4 and S11). Central Asian *P. dulcis* (PdCA) exhibited the most significant number of selective sweeps (339), followed by the wild *P. orientalis* (Po) with 224 sweeps. The two European populations, Europe I (PdE1) and Europe II (PdE2), showed consecutively lower numbers (218 and 164, respectively), and the North American population (PdNA) displayed the fewest shared sweeps. Genes under selection specifically in each cultivated almond population were associated with distinct functional categories, reflecting divergent domestication pressures. In *P. dulcis* from Central Asia, for instance, selection targeted a gene involved in cell cycle regulation, protein degradation, and floral development (e.g. CDK, encoded by gene Prudul26A028325). In *P. dulcis* from North America, the functions of candidate genes are associated with stress tolerance, cytoskeleton dynamics, and ion homeostasis, such as Prudul26A010606, which encodes a cation/H^+^ antiporter, or Prudul26A026894, which encodes a universal stress protein. We found signatures of selection in genes encoding pectinesterase (Prudul26A030900) in *P. dulcis* from Europe I or its inhibitor (Prudul26A018609) in *P. dulcis* from Central Asia. Pectinesterase has been associated with reduced juiciness, reduced levels of water soluble pectins, poor texture, and enhanced levels of insoluble pectins in peaches (Buescher & Furmanski, 1978). In *P. dulcis* from Europe II, several genes potentially involved in stress response showed positive selection signatures. For instance, Prudul26A017318 encodes a CCCH zinc finger protein, which is involved in abiotic stress responses in plants, such as salt, drought, flooding, cold temperatures, and oxidative stress (Chai et al., 2012; Han et al., 2021).

We performed a functional enrichment analysis considering the list of genes specific to each population (Figure S11) and also genes showing signatures of selection both in *P. dulcis* from Central Asia and *P. orientalis* (30 genes, Figure S11). Enriched functions in *P. dulcis* from Europe I were related to intracellular transport (Table S5). We also observed enriched function in *P. dulcis* from Central Asia that is shared with *P. orientalis*, primarily associated with the transport of sulfur compounds, and several negative regulations of metabolic processes, suggesting an increased metabolic stability that could be related to more stable environmental conditions. We also found many enriched functions in the species *P. fenzliana* (which exhibited the lowest number of sweeps, 41).

In summary, most of the genes under selection detected by at least two complementary methods were population-specific. Only a limited subset of genes overlapped across populations, with the most significant intersections involving pairwise or small group combinations, indicating that domestication and adaptation in almonds followed largely region-specific selective trajectories rather than a uniform selection signature shared across all populations.

## Discussion

By integrating genome-wide SNP data with genetic diversity estimates, population structure and demographic inferences and selection scans, we disentangled the demographic and adaptive histories of wild and cultivated almond populations. We demonstrated that *P. dulcis* maintains high genetic diversity, likely due to a weak domestication bottleneck and sustained outcrossing, while also bearing the signature of geographically structured gene flow and localized selective pressures. Central Asia emerges as a reservoir of early domesticated diversity, characterized by low levels of admixture and a high number of private alleles. In contrast, European and North American populations reflect secondary expansion fronts shaped by wild-crop introgressions, particularly from *P. orientalis*. We also identified candidate genes under selection in key metabolic pathways, including those involved in amygdalin biosynthesis and parasite resistance. Together, our findings reconstruct a multi-regional domestication landscape for almonds, an Eastern-Western diffusion history, and provide a detailed genomic roadmap for leveraging wild and cultivated diversity in future breeding programs, particularly in the context of adapting to climate change.

### A refined view of almond domestication and its consequences: An Eastern-Western diffusion shaped by gene flow

Almond belongs to the *Prunus* genus, which includes economically important fruit species such as apricot, peach, cherry, and plum (*P. domestica*) (Covert, 2011; Kester et al., 1991), yet its domestication history remains comparatively understudied. Our population genetic diversity estimates, population genetic structure, and demographic inferences suggest an eastern-to-western diffusion of the cultivated almonds. We revealed seven genetically distinct clusters: four cultivated populations of *P. dulcis* (North America, Europe I, Europe II, and Central Asia) and three wild populations (*P. fenzliana*, *P. orientalis*, and *P. spinosissima*), which confirm the strong genetic structure among cultivated almonds, supporting earlier SSR studies (Decroocq et al, 2025; Fernández I Martí et al., 2009; Halász et al., 2019) while providing greater resolution. Central Asian *P. dulcis* formed a distinct clade, separated from European and North American cultivars. Central Asian almonds retained the highest number of private alleles, suggesting an older and more diverse domestication center compared to the European and North American almonds. In the West, the two European populations were closely related and North American cultivated almonds as closely related to the two European, as reported in SSR studies (Decroocq et al., 2025) suggesting that the North American gene pool derives from European sources. The elevated and persistent LD values in the North American *P. dulcis* population are consistent with recent selection, lower effective recombination, and/or a substantial bottleneck, the latter likely associated with modern breeding and a narrow genetic base.

Our genome-wide analysis also revealed the roles of drift and gene flow in shaping the genetic diversity of cultivated almonds. Cultivated almonds retain significantly higher genetic diversity than their wild relatives, consistent with a weak domestication bottleneck typical of perennial species (Gaut et al., 2015). This lack of bottleneck can be a combination of chance seedling breeding but also wild-to-crop gene flow. While recent footprint of admixture was elusive, except among recent cultivars (mostly North American and European) and supposed-invasive New Zealander individuals in the population structure inferences, we showed patterns of historical crop-wild gene flow across the cultivated range, which suggests past crop-wild introgressions. In particular, the occurrence of gene flow between *P. orientalis* and the European and North American cultivated almonds was substantial, whereas gene flow with the Central Asian cultivated almonds was minimal, as inferred from the ABBA-BABA tests. These findings expand on previous evidence for wild-to-crop gene flow with *P. orientalis* (Delplancke et al., 2012), in contrast to earlier suggestions of limited wild-to-crop gene flow (Velasco et al., 2016). The limited gene flow with Central Asian cultivated almonds further emphasizes the isolation of the *P. spinosissima* population. Further analyses of the extent of gene flow and the genomic landscape of introgression, using additional samples of wild populations, will help pinpoint the directionality and genes involved in these introgressions.

The inclusion of wild species (*P. spinosissima*, *P. turcomanica*, *P. fenzliana,* and *P. orientalis*), which have never been studied at the genome-wide level before, provides valuable insights into the evolutionary history of almonds and raises questions about the origin of domestication. First, *P. turcomanica* grouped with *P. spinosissima* in population structure inferences but was differentiated from *P. spinosissima* in the neighbor-net; it was excluded from further analyses due to low sampling size. Our results confirmed taxonomic hypothesis on the close relatedness between *P. turcomanica* and *P. spinosissima* (Browicz & Zohary, 1996) and refined earlier SSR-based results (Decroocq et al., 2025), but further sampling will be needed to elucidate the fine-scale relationships among wild almond species and clarify the classification of poorly characterized taxa, such as *P. turcomanica*. Second, our results confirm the divergence and strong genetic differentiation between wild and cultivated populations, with species such as *P. fenzliana* and *P. spinosissima* harboring numerous private alleles and distinct genomic signatures of isolation. Second, even once recent crop-wild admixed samples were removed, the Central Asian cultivated almonds were genetically closer to the Oriental wild almond *P. orientalis* than to the Central Asian wild almonds *P. spinosissima* and *P. fenzliana*. Although a previous study suggested that *P. fenzliana* was the primary progenitor of cultivated almonds (Rahemi et al., 2012), this conclusion should be interpreted with caution. Our sampling did not cover the full diversity of wild almonds across the Middle East (including regions like Iraq, Iran, Syria), where Iranian and other researchers report additional wild species (Halász et al., 2019). Further expanded sampling will be required to reconstruct the contributions of wild taxa to cultivated almonds fully, and to determine whether *P. orientalis* played a direct progenitor role or only contributed through wild-to-crop introgression.

The New Zealand almond individuals provided another perspective on gene flow and back-to-wild evolution. Despite apparent spatial isolation, these almonds exhibited strong admixture from European and North American cultivated gene pools, and to a lesser extent from *P. spinosissima*. This pattern may result from ancestral polymorphisms retained after the introduction of almond trees ∼200 years ago (about 20 generations) by European settlers (Covert, 2011), or recent cross-pollination and gene flow among different cultivated gene pools. These almond trees grow in relatively arid environments, suggesting that their feralization process—and the observed admixture—may have favored genomic regions associated with adaptation to dry conditions. As such, New Zealand almond trees represent a unique opportunity to study feralization, local adaptation, and genetic resilience in abandoned fruit tree populations, but further sampling is now needed.

### Independent regional selection regimes during almond domestication, but common target to stress response and fruit traits

Genome-wide scans for selective sweeps, conducted using complementary methods, revealed highly heterogeneous patterns of selection across chromosomes and cultivated almond populations. There was minimal overlap in candidate genes under selection among the different cultivated populations, suggesting that domestication involved largely independent selective events across populations. However, these events often target genes involved in similar biological functions in response to stress and fruit traits.

Only a subset of 64 genes exhibiting signatures of selection were shared in at least two cultivated populations, as detected by two or more methods. These common footprints are associated with the selection of responses to stress, flowering time regulation, or kernel quality during almond domestication. Selective sweeps in TMV resistance protein N-like genes (e.g., Prudul26A000912, Prudul26A013852), previously implicated in disease resistance in potato (Hehl et al., 1999), tobacco (Marathe et al., 2002), sweet potato (Yan et al., 2024), and adzuki bean (Wang et al., 2023), suggest that disease resistance was a target during almond domestication. A gene encoding a strictosidine synthase–like protein, whose ortholog in lemon participates in alkaloid biosynthesis and contributes to domestication (Gonzalez-Ibeas et al., 2021), was under selection in all cultivated populations, suggesting a potential role in domestication-related metabolism, such as aroma production. Genes related to kernel bitterness, particularly those involved in amygdalin biosynthesis such as those encoding cytochrome P450s and UDP-glycosyltransferases, showed selection signatures, but not consistently across populations. They also did not overlap previously identified loci. This observation suggests that the evolution of sweet kernels may have involved geographically variable mechanisms or regulatory variants not captured in our scans.

We detected clear evidence that Central Asian, European, and North American populations experienced largely independent selection regimes, likely reflecting distinct environmental constraints and local breeding goals. Genes under selection specifically in each cultivated almond population were associated with various functional categories, reflecting divergent domestication pressures related to responses to stress. For instance, the cyclin-dependent kinase gene (Prudul26A028325), which is a key regulator of the cell cycle, controlling cell division and fruit development in apple (*M. domestica*), shows a selective sweep signal in *P. dulcis* from Central Asia. Prudul26A010606 in the North American almond encodes a cation/H^+^ antiporter, or Prudul26A026894, which encodes a universal stress protein. The role of cation antiporter in stress response has been observed in *M. domestica* (Mao et al., 2021). Furthermore, it has been observed that universal stress proteins have undergone gene expansion, particularly in domesticated plants like maize (*Zea mays*) and rice (*Oryza sativa*), suggesting duplication during the domestication process (Fan et al., 2024). These signatures indicate that the North American almond may have been selected for environmental adaptability under cultivation in more variable climates. We therefore reveal a complex, multi-regional domestication scenario, where independent regional domestication trajectories have left distinct genomic footprints. This is particularly remarkable because documented cases of regionally independent domestication in perennials are rare compared to annual crops. While some fruit trees, such as apricot (*Prunus armeniaca*) and lychee (*Litchi chinensis*), have undergone multiple domestication events across different geographic regions, they typically remain connected by substantial gene flow (Chen et al., 2023; Cornille et al., 2016; Delplancke et al., 2012; Diez et al., 2015; Greaves et al., 2023; Groppi et al., 2021; Hu et al., 2022).

Interestingly, European and North American populations—unlike Central Asian almonds, which appear to represent the cradle of domestication based on our results—also exhibit extensive introgressions with its wild relatives, *P. orientalis,* and share similar selective footprints, including sweeps involving genes related to pathogen resistance and environmental adaptation (including flowering time genes). One gene of particular interest is Prudul26A005915, encoding a sucrose-cleaving enzyme that provides UDP-glucose and fructose for various metabolic pathways. In *A. thaliana*, for instance, sucrose levels have been shown to influence the expression of the floral integrator gene *FT*, thereby modulating the transition from vegetative to reproductive development (Arrom & Munné-Bosch, 2012). This supports the hypothesis that variation at Prudul26A005915 in *P. orientalis* and in *P. dulcis* from Central Asia may be associated with changes in the timing or environmental responsiveness of flowering, possibly as an adaptation to local climatic conditions. Although this gene lies outside the region defined by the previously reported flowering time QTL associated with the CPSCT042 marker (∼15.5 Mb on Pd07, Sánchez-Pérez et al., 2012), its putative involvement in sugar signaling suggests that it may act in parallel or downstream of established regulators, and thus its functional role in flowering time variation deserves further investigation. However, it remains unclear whether these shared signals result from recent wild-to-crop gene flow and adaptive introgression or instead reflect the retention of ancestral polymorphism and therefore selection from standing variation. To refine these insights, further sampling will be critical to consolidate selection signals and better resolve the genomic architecture of domestication and potential adaptive introgression.

### Wild relatives as reservoirs of adaptive diversity

Wild species may be a reservoir of adaptive alleles for breeding programs. This is further corroborated by demographic inferences that revealed a shared historical bottleneck during the LGM across almond populations, followed by post-glacial expansions, which indicate that the wild species survived climate changes and harbours adaptive alleles to climate changes. This finding may suggest shared ancestral selection or wild-to-crop introgression, particularly in Europe and North America, but further investigation is needed to clarify these patterns, ideally through denser and more geographically representative sampling of *P. orientalis*. By contrast, the limited number of candidate genes detected in *P. fenzliana* and *P. spinosissima* may be attributed to both lower sampling depth and a lack of shared selective targets with cultivated almond groups. These findings highlight the importance of wild relatives as potential progenitors and reservoirs of adaptive diversity. Expanding sampling of wild relatives, particularly from underexplored regions of the Middle East and Central Asia, will be critical to fully reconstruct the domestication history of almond and leverage wild diversity for breeding resilient cultivars. In addition, our analyses highlight several promising candidate genes involved in pathogen resistance, stress response, aroma, and kernel sweetness—all key traits that are likely to have been targeted during almond domestication and diversification. Yet, even though positive selection scans are powerful tools to identify domestication-related genes, they are biased toward detecting hard sweeps and may overlook polygenic adaptation or balancing selection, which are likely to have been equally important in almond evolution. Future studies should also integrate transcriptome data, phenotypic measurements, and functional genomic approaches to validate candidate domestication genes.

## Acknowledgements

This work was supported in part by the NYU IT High Performance Computing resources, services, and staff expertise, and mostly funded by ANR JCJC PLEASURE and by a Tamkeen research fund (AD454 NYUAD) to AC. The CLAND Convergence Institute funded by ANR and INRAE funded the PhD fellowship of Celia Lougmani. About ten genotypes of almond trees were also funded by a PRIMA FREECLIMB (ANR-18-PRIM-000) project awarded to VD and IE (Grant PCI2019-103670 funded by MCIN/AEI/10.13039/501100011033 and co-financed by the European Union). We thank Naima Dlalah and Henri Duval for help in the choice and leaf sampling of INRAE GAFL almond collection.

## Data availability

All scripts used for the analyses presented in this study are available at the Github link https://github.com/CornilleEclecticLab/Almond-SNP. VCF files are available on https://doi.org/10.5281/zenodo.15572161.

## Authors contributions

AC, VD, and KA obtained funding and designed the experiment. AV, CL, AM, VD, QTB, SL, DC, IE, and AC prepared the material; CL, AM, RF, XC and AC analyzed the data; All co-authors discussed the results. CL, AM, RF, AC wrote the manuscript with critical input from other co-authors.

## Funding

This research was funded by ANR JCJC PLEASURE ANR-21-CE20-0005 and Tamkeen research fund under AD454 NYUAD led by AC. The PhD fellowship of Celia Lougmani supervised by AC, VD and KA was funded by CLAND Convergence Institute funded by ANR and INRAE. About ten genotypes of almond trees were funded by a PRIMA FREECLIMB (ANR-18-PRIM-000) project awarded to VD and IE (Grant PCI2019-103670 funded by MCIN/AEI/10.13039/501100011033 and co-financed by the European Union).

## Conflict of interest

The authors of this preprint declare that they have no financial conflict of interest with the content of this article.

## Notes

### Competing Interest Statement

The authors have declared no competing interest.

