## Supplementary tables and figures for "Genomic footprints of domestication in almond (*Prunus dulcis*)"

**Table S1. Summary of almond accessions, sequencing information, and genome mapping statistics.**

This table provides a comprehensive overview of the almond accessions used in this study, along with the peach (*Prunus persica*) outgroup samples. For each accession, we report species identity, population assignment, geographic origin, sequencing platform, read length, sequencing depth, and mapping statistics against the *Prunus dulcis* cv. Texas v2.0 reference genome. Summary metrics include average genome coverage, percentage of mapped reads, and percentage of properly paired reads. This dataset was used as the foundation for all downstream analyses of population structure, demographic inference, and selection scans.

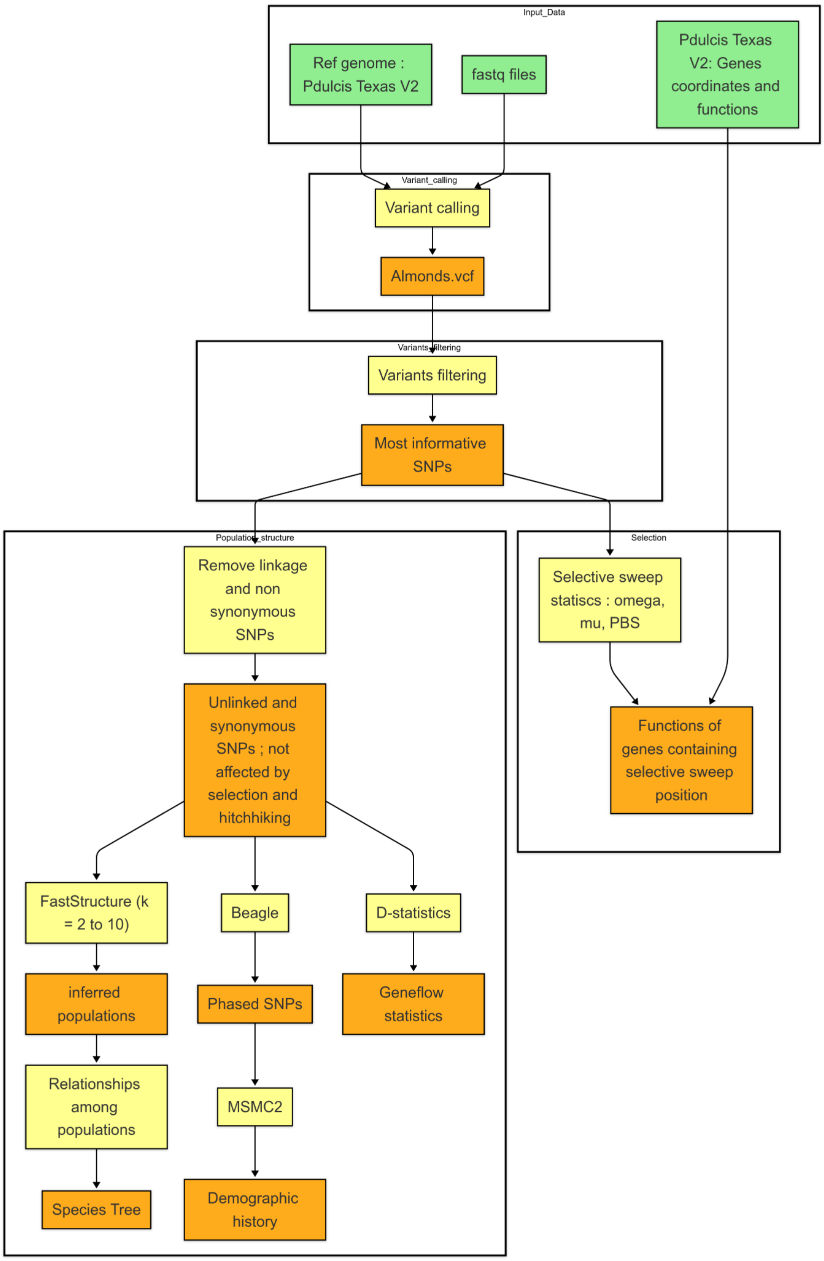


**Figure S1. Genomic analysis workflow for variant discovery and population genomics in almond.** Schematic representation of the bioinformatic pipeline used to process whole-genome resequencing data from *Prunus dulcis* and related wild species. Raw sequencing reads (FASTQ files) were aligned to the *P. dulcis* cv. Texas v2.0 reference genome, using corresponding gene annotations for downstream analyses. Variant calling and stringent filtering produced a high-confidence SNP dataset. For population genomic analyses—including structure, demographic inferences, and introgression—only unlinked synonymous SNPs were retained to reduce biases associated with linkage and selection. Population structure was inferred with *fast*STRUCTURE (K=2–10), and demographic histories were reconstructed using MSMC2 after haplotype phasing with BEAGLE. Wild-crop gene flow occurrence was quantified using ABBA-BABA D-statistics (Dsuite). Genome-wide scans for positive selection were performed using three complementary approaches: RAiSD (μ statistic), OmegaPlus (ω statistic), and the Population Branch Statistic (PBS), targeting both hard and soft sweeps as well as lineage-specific selection. Genomic regions under selection were annotated using the *P. dulcis* reference gene models to identify putative domestication and adaptation genes.

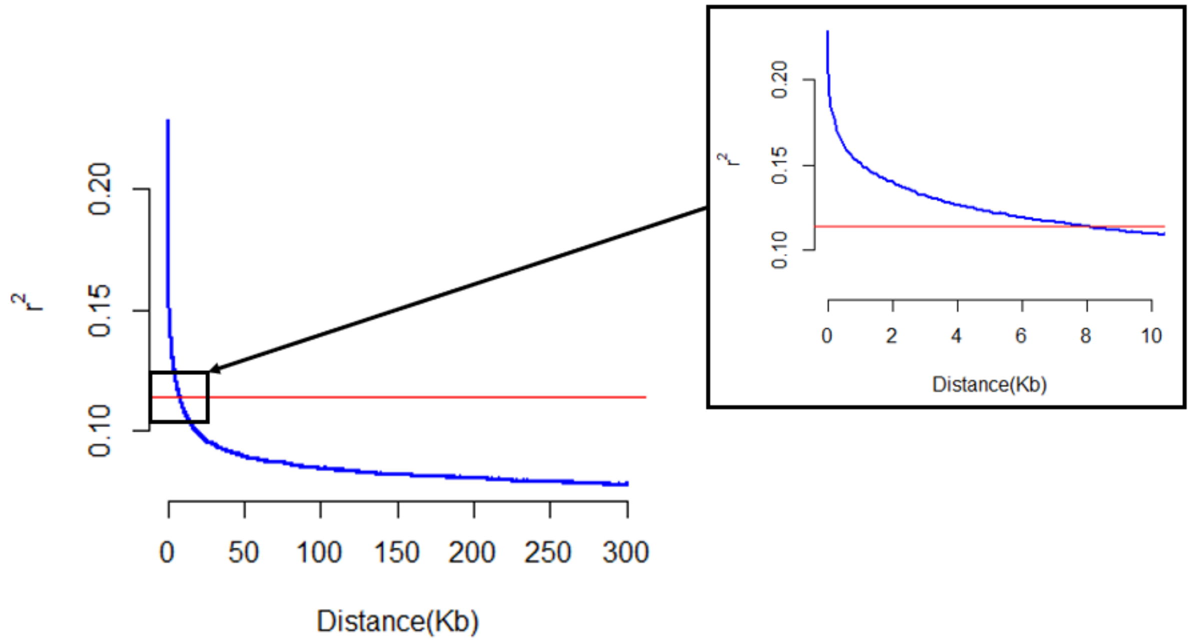


**Figure S2**. **Linkage disequilibrium (LD) decay across almond genomes.** Decay of LD measured as the squared correlation coefficient ($r^2$) between SNP pairs plotted against physical distance (kilobases, Kb), calculated using PopLDdecay. The blue curve represents the genome-wide average LD decay across all almond populations. The red horizontal line marks the point at which $r^2$ falls to half of its maximum value, used here to define the LD decay threshold. The inset panel (right) provides a magnified view of the initial segment of the decay curve, where LD declines most rapidly. LD decayed to half its maximum value at approximately 8 Kb, indicating that SNPs within this distance were considered linked and excluded from downstream analyses requiring independence (e.g., population structure and diversity estimates).

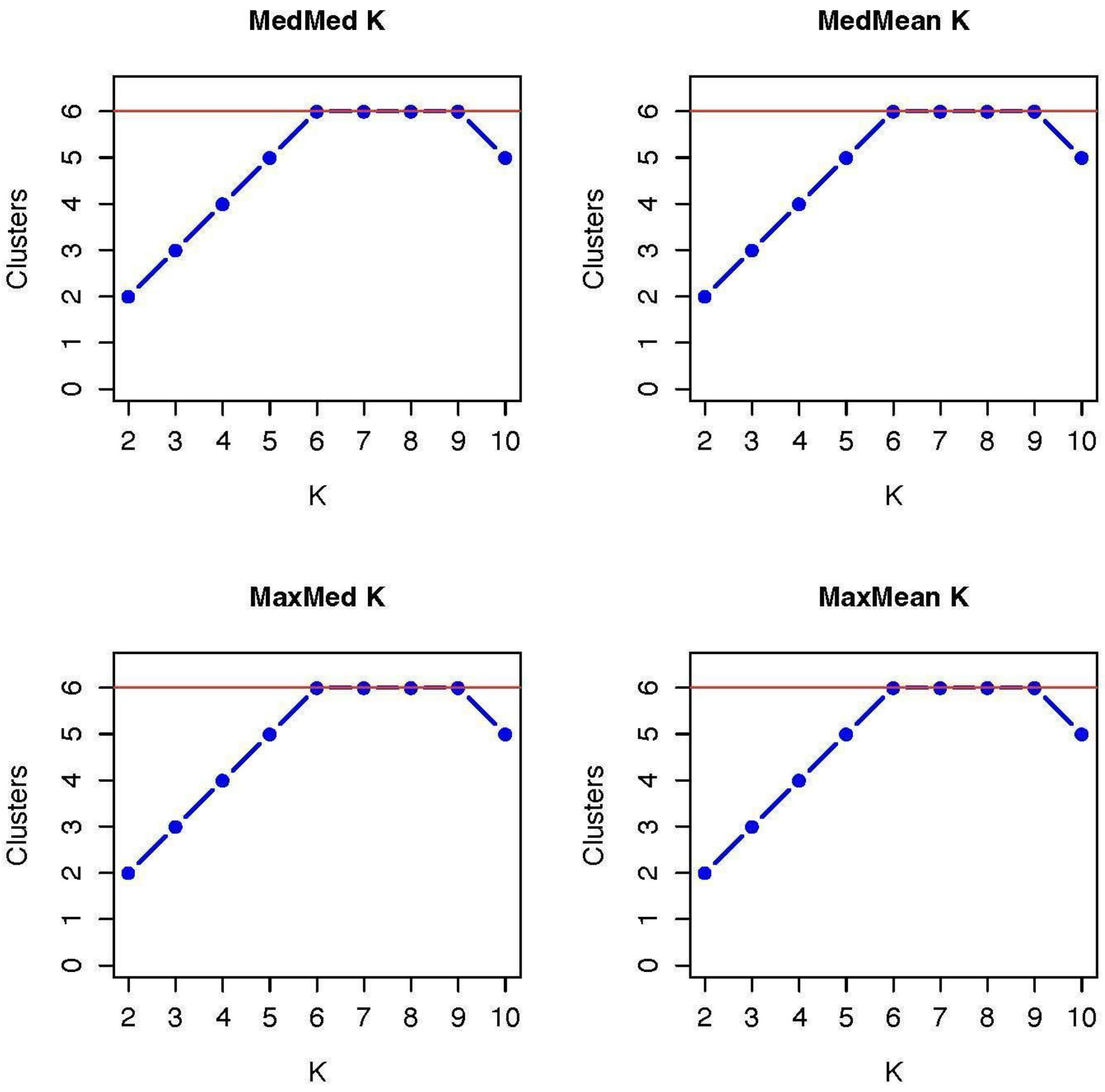


**Figure S3.** Optimal number of genetic clusters (K) estimated using summary statistics of Q-matrices from fastStructure. Plots show the optimal number of inferred clusters (K) based on four summary statistics applied to Q-matrices: MedMed, MedMean, MaxMed, and MaxMean. Each curve represents one aggregation method, with the x-axis indicating the tested K values (from 2 to 10) and the y-axis representing the corresponding statistic. The red horizontal lines mark the optimal K values estimated by each method. All four approaches converge on a range of K = 6 to 9, suggesting hierarchical population structure among almond accessions. These results support the presence of fine-scale differentiation within both wild and cultivated *Prunus* populations, in line with barplot and PCA-based structure analyses.

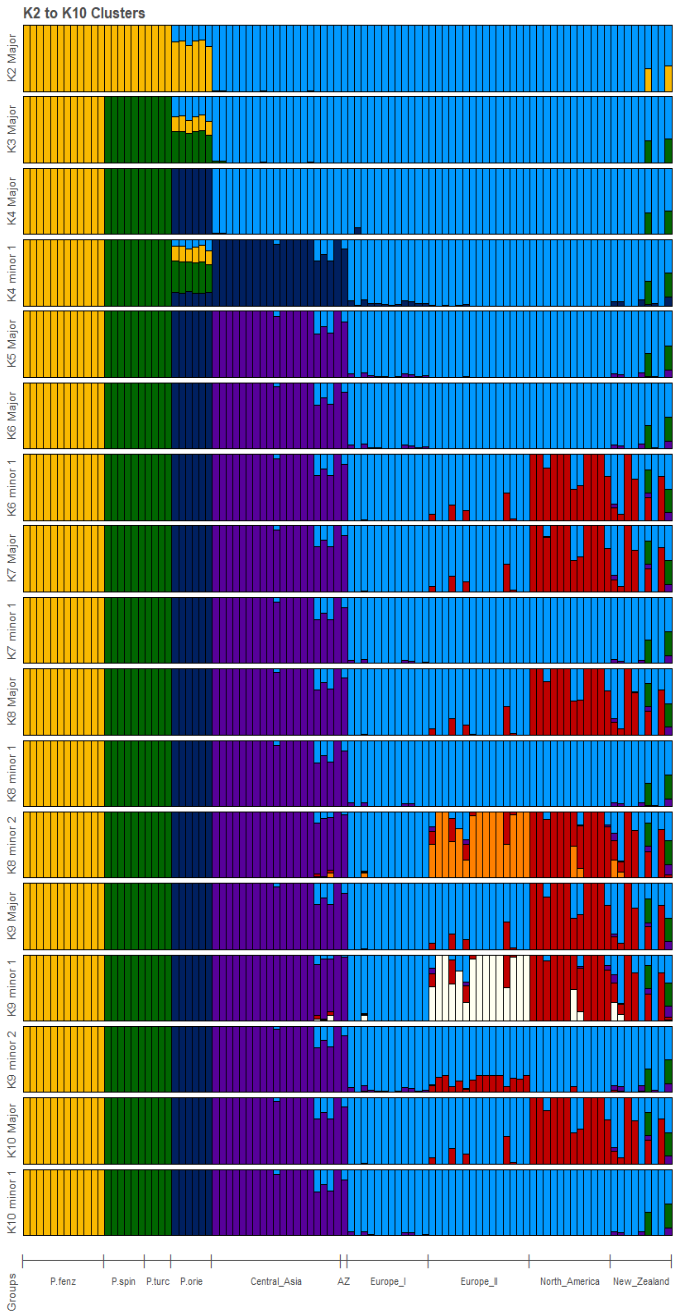


**Figure S4. Hierarchical population structure of wild and cultivated almonds inferred using fastStructure (K = 2–10).** Bar plots display individual ancestry proportions estimated with fastStructure for values of K ranging from 2 to 10, based on 13,167 unlinked synonymous SNPs. Each vertical bar represents one individual, and the colored segments indicate the proportion of the genome assigned to each of the inferred genetic clusters. Samples are grouped along the x-axis by species (wild: *P. fenzliana* [P. fenz], *P. spinosissima* [P. spin], *P. turcomanica* [P. turc], *P. orientalis* [P. orie]) and cultivated population origin (*P. dulcis* from Central Asia, Azerbaijan [AZ], Europe I, Europe II, North America, and New Zealand). As K increases, deeper levels of population substructure become evident. At low K, the major distinction between wild and cultivated almonds emerges, while higher K values reveal finer-scale differentiation among cultivated groups, especially between Central Asian, European, and North American P. dulcis. Admixed individuals, particularly from New Zealand, display complex ancestry profiles derived from multiple cultivated lineages.

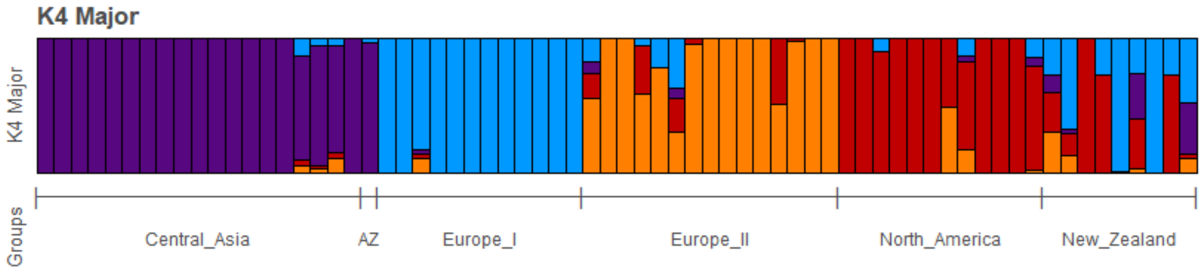


**Figure S5. Population structure of 68 cultivated almonds inferred using *fast*STRUCTURE for K = 4.** Each vertical bar represents an individual genotype, partitioned into colored segments that correspond to the individual's estimated membership proportion in each of the four inferred ancestral clusters. Individuals are grouped by geographic origin: Central Asia, Azerbaijan (AZ), Europe I, Europe II, North America, and New Zealand.

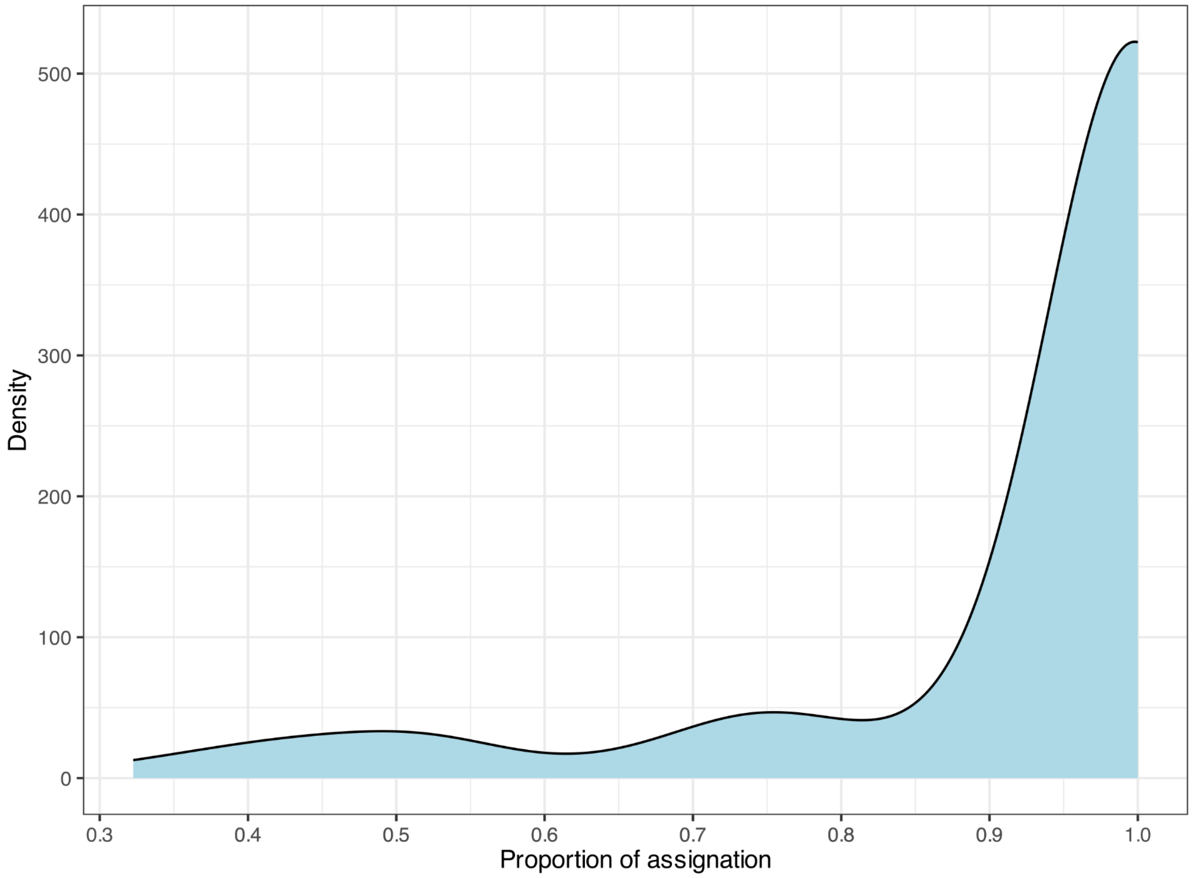


**Figure S6. Density distribution of individual ancestry coefficients across all inferred genetic clusters.** The x-axis represents the maximum proportion of genome assignment of each individual to a single cluster (Q value), as inferred by *fast*STRUCTURE. Most individuals show high assignment probabilities (Q > 0.85), indicating strong genetic clustering, while a smaller subset displays intermediate values, suggestive of potential admixture or recent gene flow or ancestral polymorphism.

**Table S2. Membership coefficients** (Q-values) **from** for each individual almond accession based on *fast*STRUCTURE analyses at $K$ = 8**.** Columns representing the proportion of its genome assigned to each of the eight inferred genetic clusters. Accessions with membership coefficients >0.85 in a single cluster are considered confidently assigned to that genetic group, whereas individuals with Q-values <0.85 across multiple clusters are classified as admixed. These coefficients were used to define population assignments for downstream analyses of genetic diversity, demographic history, and selection.

| sample ID | Blue cluster | Yellow cluster | Green cluster | Dark blue cluster | Purple cluster | Red cluster | Orange Cluster | Coef. Pop8 | Maximum assignation coefficient |
|---|---|---|---|---|---|---|---|---|---|
| BBQ1155 | 0.3226 | 0 | 0 | 0 | 0.1165 | 0.2916 | 0.2693 | 0 | 0.3226 |
| BBQ1323 | 0.746 | 0 | 0 | 0 | 0.0058 | 0.1532 | 0.095 | 0 | 0.746 |
| BBQ1325 | 0 | 0 | 0 | 0 | 0 | 1 | 0 | 0 | 1 |
| BBQ1335 | 0.2851 | 0 | 0 | 0 | 0 | 0.7149 | 0 | 0 | 0.7149 |
| BBQ1339 | 1 | 0 | 0 | 0 | 0 | 0 | 0 | 0 | 1 |
| BBQ1347 | 0.1653 | 0 | 0.3513 | 0 | 0.0888 | 0.3946 | 0 | 0 | 0.3946 |
| BBQ1351 | 1 | 0 | 0 | 0 | 0 | 0 | 0 | 0 | 1 |
| BBQ1353 | 0.2728 | 0 | 0 | 0 | 0 | 0.7272 | 0 | 0 | 0.7272 |
| BBQ1355 | 0.4341 | 0 | 0.359 | 0 | 0.1599 | 0.0456 | 0.0014 | 0 | 0.4341 |
| IBO3401 | 0 | 0 | 0 | 0 | 0 | 1 | 0 | 0 | 1 |
| IBO3402 | 0.2232 | 0 | 0 | 0 | 0.0726 | 0.1897 | 0.5144 | 0 | 0.5144 |
| IBO3403 | 0 | 0 | 0 | 0 | 0 | 0 | 1 | 0 | 1 |
| IBO3404 | 0 | 0 | 0 | 0 | 0 | 0 | 1 | 0 | 1 |
| IBO3405 | 1 | 0 | 0 | 0 | 0 | 0 | 0 | 0 | 1 |
| IBO3406 | 1 | 0 | 0 | 0 | 0 | 0 | 0 | 0 | 1 |
| IBO3407 | 0 | 0 | 0 | 0 | 0 | 1 | 0 | 0 | 1 |
| IBO3408 | 0.099 | 0 | 0 | 0 | 0 | 0.3561 | 0.5448 | 0 | 0.5448 |
| IBO3409 | 0.251 | 0 | 0 | 0 | 0 | 0 | 0.749 | 0 | 0.749 |
| IBO3410 | 0.4239 | 0 | 0 | 0 | 0.0629 | 0.2471 | 0.266 | 0 | 0.4239 |
| IBO3411 | 0.9023 | 0 | 0 | 0 | 0.0107 | 0.0154 | 0.0716 | 0 | 0.9023 |
| ORI3087 | 0 | 0 | 0 | 1 | 0 | 0 | 0 | 0 | 1 |
| ORI3088 | 0 | 0 | 0 | 1 | 0 | 0 | 0 | 0 | 1 |
| ORI3089 | 0 | 0 | 0 | 1 | 0 | 0 | 0 | 0 | 1 |
| ORI3091 | 0 | 0 | 1 | 0 | 0 | 0 | 0 | 0 | 1 |
| ORI3092 | 0 | 0 | 1 | 0 | 0 | 0 | 0 | 0 | 1 |
| ORI3093 | 0 | 0 | 1 | 0 | 0 | 0 | 0 | 0 | 1 |
| ORI3094 | 0 | 0 | 1 | 0 | 0 | 0 | 0 | 0 | 1 |
| ORI3112 | 0 | 0 | 0 | 1 | 0 | 0 | 0 | 0 | 1 |
| ORI3113 | 0 | 0 | 0 | 1 | 0 | 0 | 0 | 0 | 1 |
| ORI3114 | 0 | 0 | 0 | 1 | 0 | 0 | 0 | 0 | 1 |
| SRR1032 | 0 | 0 | 0 | 0 | 1 | 0 | 0 | 0 | 1 |
| SRR1040 | 0 | 0 | 0 | 0 | 1 | 0 | 0 | 0 | 1 |
| SRR1049 | 0 | 0 | 0 | 0 | 1 | 0 | 0 | 0 | 1 |
| SRR1057 | 0 | 0 | 0 | 0 | 1 | 0 | 0 | 0 | 1 |
| SRR1065 | 0 | 0 | 0 | 0 | 1 | 0 | 0 | 0 | 1 |
| SRR1073 | 0 | 0 | 0 | 0 | 1 | 0 | 0 | 0 | 1 |
| SRR1083 | 0 | 0 | 0 | 0 | 1 | 0 | 0 | 0 | 1 |
| SRR1098 | 0 | 0 | 0 | 0 | 1 | 0 | 0 | 0 | 1 |
| SRR1113 | 0 | 0 | 0 | 0 | 1 | 0 | 0 | 0 | 1 |
| SRR1181 | 0 | 0 | 0 | 0 | 1 | 0 | 0 | 0 | 1 |

| | | | | | | | | | |
|---|---|---|---|---|---|---|---|---|---|
| SRR1192 | 0 | 0 | 0 | 0 | 1 | 0 | 0 | 0 | 1 |
| SRR1204 | 0 | 0 | 0 | 0 | 1 | 0 | 0 | 0 | 1 |
| SRR1229 | 0.1143 | 0 | 0 | 0 | 0 | 0.8857 | 0 | 0 | 0.8857 |
| SRR1238 | 0 | 0 | 0 | 0 | 0 | 1 | 0 | 0 | 1 |
| VDS1301 | 0 | 0 | 0 | 0 | 0 | 1 | 0 | 0 | 1 |
| VDS1302 | 0 | 0 | 0 | 0 | 0 | 0.0638 | 0.9362 | 0 | 0.9362 |
| VDS1303 | 1 | 0 | 0 | 0 | 0 | 0 | 0 | 0 | 1 |
| VDS1304 | 0 | 1 | 0 | 0 | 0 | 0 | 0 | 0 | 1 |
| VDS1305 | 0 | 0 | 0 | 0 | 1 | 0 | 0 | 0 | 1 |
| VDS1306 | 0 | 0 | 1 | 0 | 0 | 0 | 0 | 0 | 1 |
| VDS1307 | 0 | 1 | 0 | 0 | 0 | 0 | 0 | 0 | 1 |
| VDS1308 | 0 | 0 | 1 | 0 | 0 | 0 | 0 | 0 | 1 |
| VDS1309 | 0 | 0 | 0 | 0 | 0 | 0 | 1 | 0 | 1 |
| VDS1310 | 1 | 0 | 0 | 0 | 0 | 0 | 0 | 0 | 1 |
| VDS1312 | 0 | 0 | 0 | 0 | 1 | 0 | 0 | 0 | 1 |
| VDS1313 | 0 | 0 | 0 | 0 | 0 | 0 | 1 | 0 | 1 |
| VDS1314 | 0 | 1 | 0 | 0 | 0 | 0 | 0 | 0 | 1 |
| VDS1315 | 0 | 0 | 1 | 0 | 0 | 0 | 0 | 0 | 1 |
| VDS1316 | 0 | 0 | 0 | 0 | 0 | 0 | 1 | 0 | 1 |
| VDS1317 | 0 | 0 | 0 | 0 | 1 | 0 | 0 | 0 | 1 |
| VDS1318 | 0 | 0 | 1 | 0 | 0 | 0 | 0 | 0 | 1 |
| VDS1319 | 0 | 1 | 0 | 0 | 0 | 0 | 0 | 0 | 1 |
| VDS1320 | 0 | 0 | 0 | 0 | 0 | 0 | 1 | 0 | 1 |
| VDS1321 | 0 | 0 | 1 | 0 | 0 | 0 | 0 | 0 | 1 |
| VDS1322 | 1 | 0 | 0 | 0 | 0 | 0 | 0 | 0 | 1 |
| VDS1323 | 0.1655 | 0 | 0 | 0 | 0.7742 | 0.0475 | 0.0129 | 0 | 0.7742 |
| VDS1324 | 0 | 0 | 0 | 0 | 0 | 0.4968 | 0.5032 | 0 | 0.5032 |
| VDS1325 | 0 | 0 | 0 | 0 | 0 | 0.0375 | 0.9625 | 0 | 0.9625 |
| VDS1326 | 0 | 1 | 0 | 0 | 0 | 0 | 0 | 0 | 1 |
| VDS1328 | 0 | 1 | 0 | 0 | 0 | 0 | 0 | 0 | 1 |
| VDS1329 | 0.0962 | 0 | 0 | 0 | 0.8889 | 0.0149 | 0 | 0 | 0.8889 |
| VDS1330 | 0 | 0 | 0 | 0 | 0 | 1 | 0 | 0 | 1 |
| VDS1331 | 1 | 0 | 0 | 0 | 0 | 0 | 0 | 0 | 1 |
| VDS1332 | 0 | 1 | 0 | 0 | 0 | 0 | 0 | 0 | 1 |
| VDS1333 | 0 | 1 | 0 | 0 | 0 | 0 | 0 | 0 | 1 |
| VDS1382 | 0 | 0 | 0 | 0 | 0 | 0 | 1 | 0 | 1 |
| VDS1383 | 1 | 0 | 0 | 0 | 0 | 0 | 0 | 0 | 1 |
| VDS1384 | 0.0862 | 0 | 0 | 0 | 0.7937 | 0.0439 | 0.0763 | 0 | 0.7937 |
| VDS1385 | 1 | 0 | 0 | 0 | 0 | 0 | 0 | 0 | 1 |
| VDS1386 | 0 | 0 | 0 | 0 | 1 | 0 | 0 | 0 | 1 |
| VDS1387 | 0 | 1 | 0 | 0 | 0 | 0 | 0 | 0 | 1 |
| VDS1388 | 0 | 0 | 1 | 0 | 0 | 0 | 0 | 0 | 1 |
| VDS1389 | 1 | 0 | 0 | 0 | 0 | 0 | 0 | 0 | 1 |
| VDS1390 | 0 | 0 | 0 | 0 | 0 | 0.5187 | 0.4813 | 0 | 0.5187 |
| VDS1391 | 0 | 1 | 0 | 0 | 0 | 0 | 0 | 0 | 1 |
| VDS1392 | 0.1927 | 0 | 0 | 0 | 0.0135 | 0.6564 | 0.1374 | 0 | 0.6564 |
| VDS1393 | 1 | 0 | 0 | 0 | 0 | 0 | 0 | 0 | 1 |
| VDS1394 | 0 | 0 | 0 | 0 | 0 | 1 | 0 | 0 | 1 |
| VDS1396 | 0.0438 | 0 | 0 | 0 | 0.9561 | 0 | 0 | 0 | 0.9561 |
| VDS1401 | 1 | 0 | 0 | 0 | 0 | 0 | 0 | 0 | 1 |

| | | | | | | | | | |
|---|---|---|---|---|---|---|---|---|---|
| VDS1402 | 0 | 0 | 0 | 0 | 0 | 0 | 1 | 0 | 1 |
| VDS1403 | 0 | 0 | 0 | 0 | 0 | 1 | 0 | 0 | 1 |
| VDS1404 | 0 | 0 | 0 | 0 | 0 | 1 | 0 | 0 | 1 |
| VDS1405 | 0.1998 | 0 | 0 | 0 | 0.0323 | 0.7678 | 0 | 0 | 0.7678 |
| VDS1406 | 0 | 1 | 0 | 0 | 0 | 0 | 0 | 0 | 1 |
| VDS1407 | 0 | 1 | 0 | 0 | 0 | 0 | 0 | 0 | 1 |

**Table S3. Pairwise comparisons of nucleotide diversity (π) between populations using the Mann–Whitney U test.** The values correspond to p-values from pairwise Mann–Whitney U tests comparing nucleotide diversity (π) between populations. P-values equal to 0 indicate values below machine precision (P < 1e−300).

| | *P. dulcis* Central Asia | *P. dulcis* North America | *P. dulcis* Europe II | *P. dulcis* Europe I | *P. orientalis* | *P. fenzliana* | *P. spinosissima* |
|---|---|---|---|---|---|---|---|
| *P. dulcis* Central Asia | - | | | | | | |
| *P. dulcis* North America | 4.67 e−288 | - | | | | | |
| *P. dulcis* Europe II | 1.99 e−69 | 1.11 e−79 | - | | | | |
| *P. dulcis* Europe I | 2.15 e−10 | 2.76 e−193 | 1.69 e−28 | - | | | |
| *P. orientalis* | 0 | 0 | 0 | 0 | - | | |
| *P. fenzliana* | 0 | 8.81 e−99 | 0 | 0 | 0 | - | |

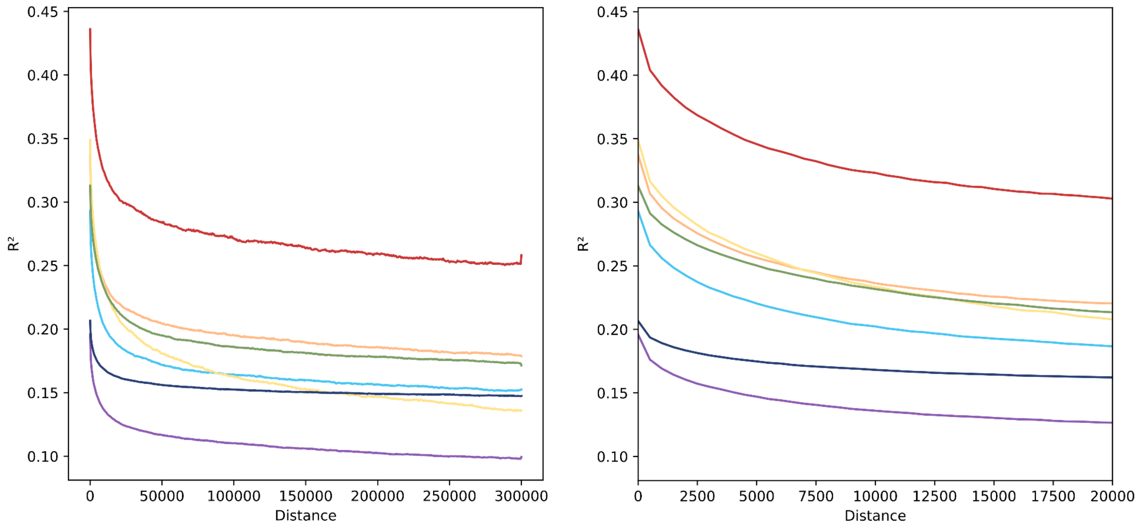


**Figure S7. Linkage disequilibrium (LD) decay with physical distance between SNP pairs in seven almond populations (i.e., group of individuals assigned with a membership coefficient >0.85 inferred at *K*=8 with fastSTRUCTURE).** The left panel shows genome-wide LD decay up to 300 kb, while the right panel zooms in on the first 20 kb, where LD decays most steeply. Each curve represents a distinct population: *Prunus dulcis* from Central Asia, purple, European cultivated populations in light and dark blue, North American population in red; *P. fenzliana* in yellow, *P. orientalis* in navy; *P. spinosissima* in green. The North American population displays the highest initial LD values and the slowest decay, indicating extended haplotype blocks possibly resulting from recent selective sweeps, bottlenecks, or reduced recombination. In contrast, wild species (*P. fenzliana*, *P. orientalis*, *P. spinosissima*) show comparatively rapid LD decay, reflecting long-term recombination dynamics and smaller historical effective linkage blocks, despite their lower current nucleotide diversity.

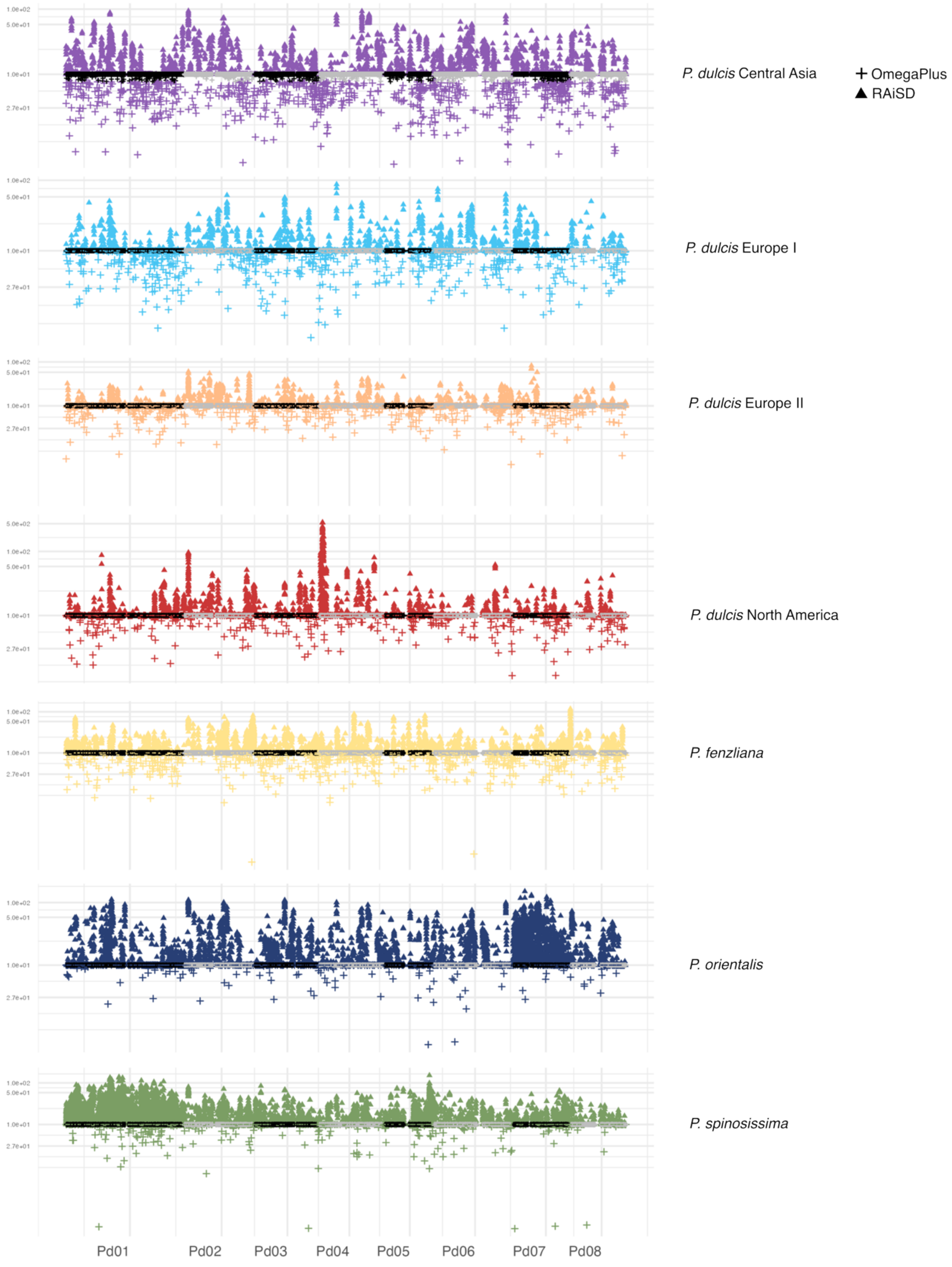


**Figure S8. Genome-wide detection of selective sweeps in cultivated and wild almond populations** (6,063,769 SNPs). Selection signal analysis was performed on the whole genome of cultivated and wild almond populations using OmegaPlus and RAiSD methods. The curves represent the statistics associated with each method, with peaks corresponding to candidate regions for selective sweep. Outlier thresholds are indicated by horizontal lines, thus allowing the identification of regions with significant deviation from the genomic background.

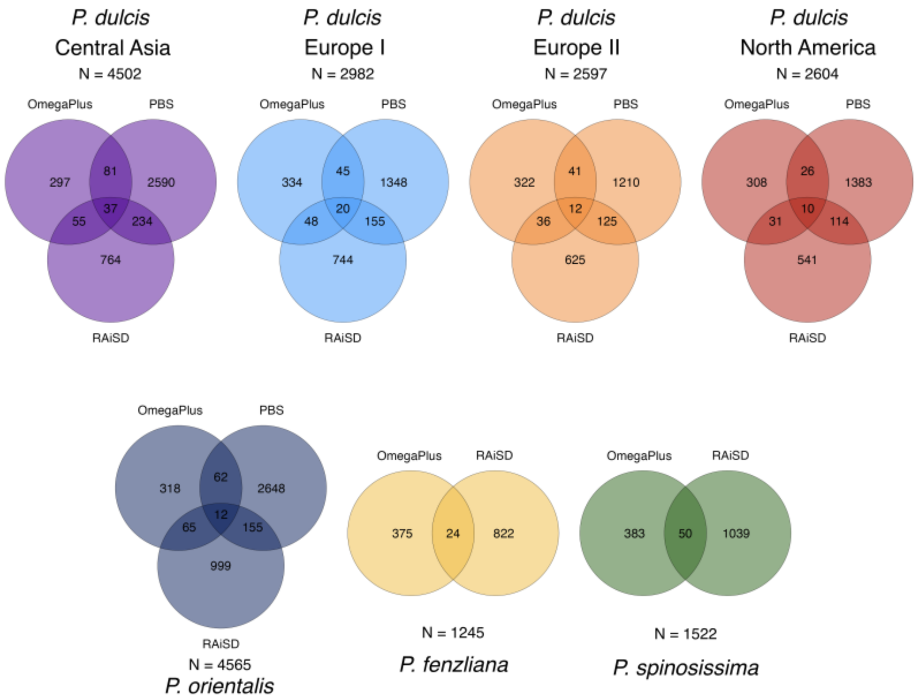


**Figure S9. Overlap of genes under selection detected by three complementary methods across almond populations.** Venn diagrams showing the number of genes identified as candidates for selective sweeps using three different methods: RAiSD, OmegaPlus, and PBS (Population Branch Statistic). Results are shown for *P. dulcis* populations from Central Asia, Europe I, Europe II, and North America, as well as for the wild species *P. orientalis*, *P. fenzliana*, and *P. spinosissima*. For *P. fenzliana* and *P. spinosissima*, only RAiSD and OmegaPlus were applied. The total number of candidate genes per population is indicated above each diagram. The overlaps illustrate the consistency and complementarity of the methods used to detect selective sweep signatures.

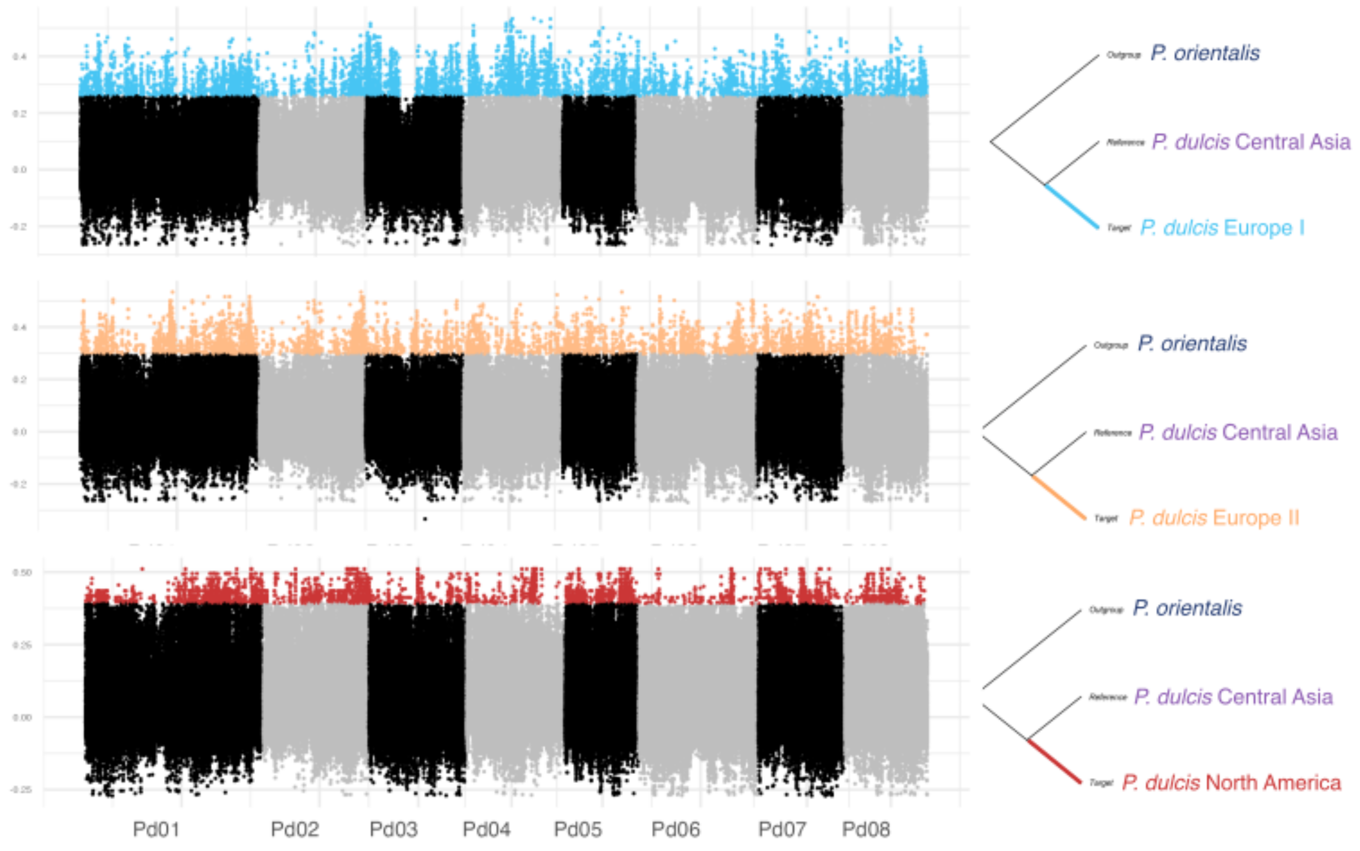


**Figure S10.** Genome-wide detection of population-specific selective sweeps in cultivated almond populations using PBS (common 744,241 SNPs among populations). PBS (Population Branch Statistic) values were calculated genome-wide for three cultivated *Prunus dulcis* populations (Europe I, Europe II, and North America), using *P. dulcis* Central Asia as the reference and *P. orientalis* as the outgroup in each case. The x-axis represents the eight pseudomolecules (Pd01–Pd08) of the almond genome, and the y-axis shows the PBS value per genomic window. Colored dots highlight windows with elevated PBS values specific to the target population (Europe I: blue, Europe II: orange, North America: red), indicating putative regions under recent positive selection, while Grey and black dots represent positions whose score is below the 1% cutoff. Tree topologies used for each PBS comparison are shown on the right.

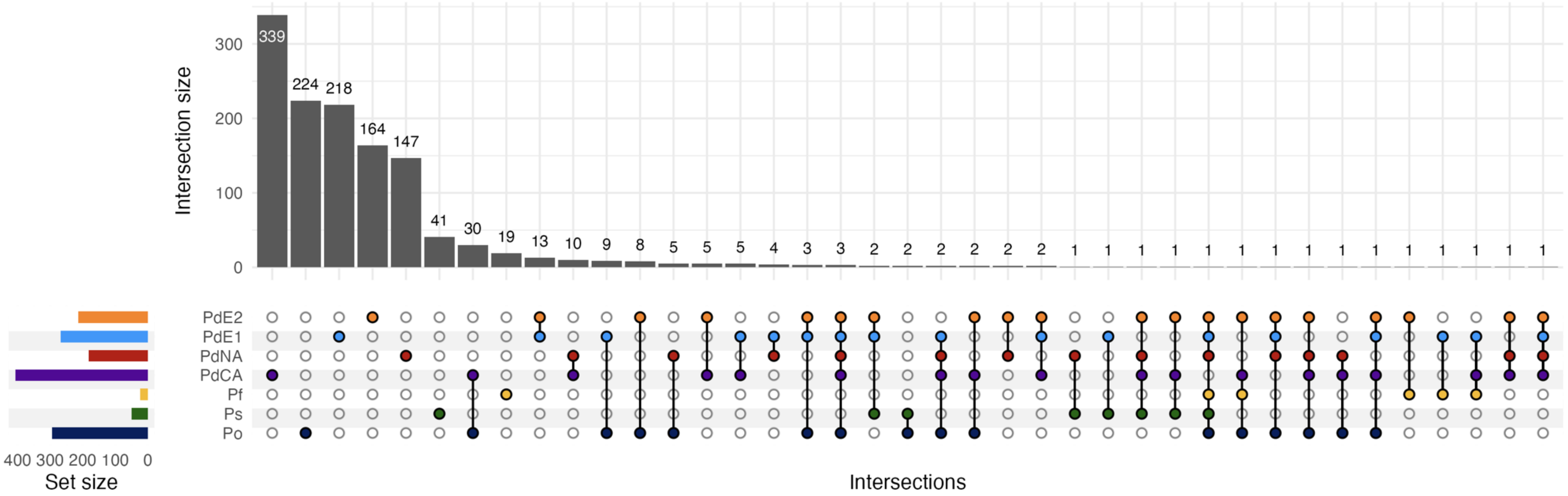


**Figure S11. Shared and population-specific genes under selection across almond populations.** Upset plot showing the number of overlapping genes carrying signatures of selective sweeps identified by at least two methods (OmegaPlus and/or RAiSD and/or PBS) across the seven almond populations analyzed. Each bar on the top panel represents the number of genes shared among the populations indicated by the connected dots in the matrix below. The left panel shows the total number of genes detected per population. Populations are coded by color: *P. dulcis* Central Asia (PdCA), Europe I (PdE1), Europe II (PdE2), North America (PdNA), *P. fenzliana* (Pf), *P. orientalis* (Po), and *P. spinosissima* (Ps). Most genes under selection are population-specific, but several overlaps indicate shared selection signals across cultivated and/or wild populations, possibly reflecting common adaptive or domestication-related targets.

**Table S4**. List of genes containing selective sweep signatures detected using at least two methods among RAiSD, OmegaPlus and PBS, in wild and cultivated almonds (excel file Table_S4.xlsx).

**Table S5**. Go term enriched in genes containing selective sweep signatures in almonds populations (excel file Table_S5.xlsx)

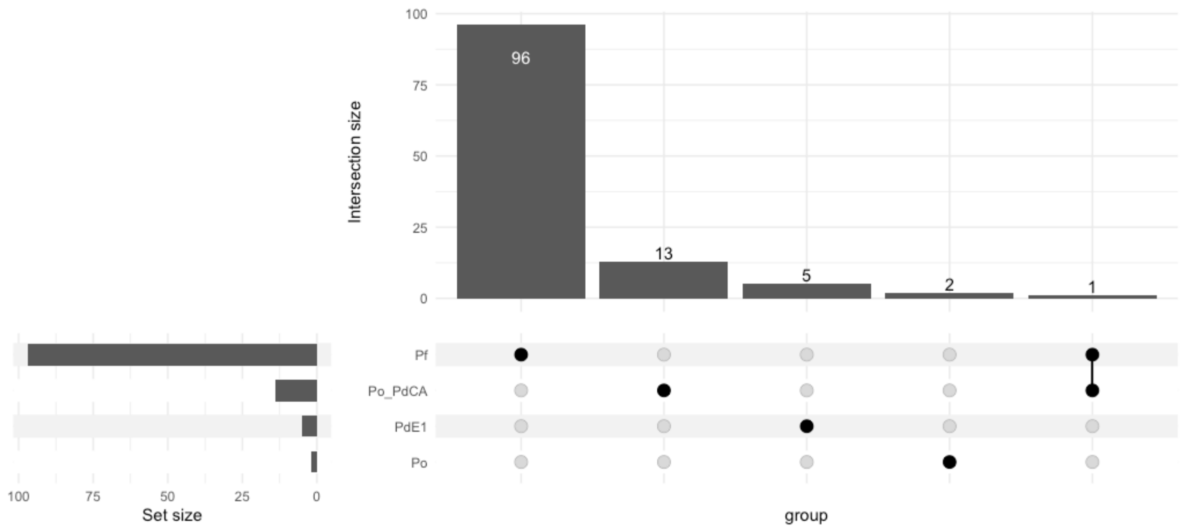


**Figure S12. Shared and specific enriched functions among gene sets under selection in different populations.** The UpSet plot shows the intersection of enriched GO terms identified in gene sets specific to three populations: PdCA (*P. dulcis* from Central Asia), PdE1 (*P. dulcis* from Europe I), Pf (*P. fenzliana*, wild), and a shared gene set between PdCA and Po (*P. orientalis*, putative progenitor). Each bar represents the number of enriched GO terms specific to a population or shared between groups, as indicated by the connected dots below. The left panel indicates the total number of enriched terms per population. The majority of functions are unique to individual populations, 13 GO terms are shared between PdCA and Po, suggesting potential inheritance or convergent selection.
